# Systems-level Consequences of Low RAF Abundance for EGFR-ERK Signaling

**DOI:** 10.1101/2025.06.12.659034

**Authors:** Sung Hyun Lee, Paul J. Myers, Kevin S. Brown, Leslie M. Loew, Alexander Sorkin, Matthew J. Lazzara

**Author notes:** Corresponding Author: Matthew J. Lazzara 102 Engineers’ Way, PO Box 400741 Charlottesville, Virginia 22904-4741. Grant support: NSF MCB 1716537, NSF MCB 1715132, NSF MCB 1716075, NSF MCB 1715342, NIH U54 CA274499, NIH R35 GM148363, NIH T32LM012416, NIH R24 GM137787. Potential conflicts of interest: The authors declare no conflicts of interest.

## Abstract

The RAF kinases are central links between RAS, once activated by receptor tyrosine kinases (RTKs), and the extracellular signal-regulated kinases (ERK). In many cancer cells, RAFs are the least abundantly expressed RTK-ERK pathway proteins and can be present at just hundreds of copies per cell at the plasma membrane, but the consequences of limited RAF expression are unclear. By developing continuum and stochastic computational models of the epidermal growth factor receptor (EGFR)-ERK pathway, we showed that low RAF abundance creates stoichiometric bottlenecks between RTKs and ERK with concomitant stochastic RAF dynamics that propagate to weakly expressed downstream pathway proteins. Advanced sensitivity and Sloppiness analyses identified RAS activation and RAS-RAF interactions as strong determinants of signaling in low-RAF settings and revealed an efficient model fitting approach. RAF bottlenecks were predicted to impede ERK activation by oncogenic RAS mutants and explained a tendency for RAF1 membrane localization to be noisy. This work provides quantitative insight into a common, yet unexplored, regime for EGFR-ERK signaling and a systematic approach to develop and characterize dynamic models of receptor-mediated signaling.

**STATEMENT OF SIGNIFICANCE:** RAF kinases connect receptors to the mitogenic ERK signaling pathway by translocating to the plasma membrane, but in a substantial fraction of cancer cell contexts RAFs are greatly outnumbered by other pathway proteins, potentially creating an unrecognized and consequential signaling bottleneck. We trained a novel computational model of EGFR-ERK signaling and characterized it comprehensively using integrated multivariate sensitivity analyses and Sloppiness analysis. The results revealed that low RAF abundance suppresses EGFR-mediated ERK activation, limits the effects of upstream oncogenic RAS mutants, and creates stochastic RAF dynamics that can propagate downstream. Thus, the canonical EGFR-ERK pathway exhibits divergent behaviors in a parameter space representative of a substantial fraction of cancer cell settings.

## INTRODUCTION

The EGFR-ERK signaling pathway is one of the most extensively studied biochemical networks due to its ubiquitous roles in normal cell physiology and disease (1). While the structural and functional properties of the constituent proteins are well characterized (2), it is only recently that mass spectrometry enabled absolute quantification of protein kinases and functional protein complexes in the pathway (3). We now understand that some pathway nodes and protein complexes exist in surprisingly low abundance (<1000 copies/cell), suggesting potential rate limitations in signal transduction. For example, the multifunctional adapter protein GRB2-associated binding protein 1 (GAB1) is expressed at ∼1500 copies/cell in HeLa (4), limiting its ability to activate the abundant protein tyrosine phosphatase SHP2 (∼300,000 copies/cell) by forming the GAB1:SHP2 complexes that regulate signaling downstream of EGFR and other receptor tyrosine kinases (RTKs) (5,6). Signal-initiating receptors themselves can be less abundant than their common adapters. For example, with less than half of EGFR tyrosine-phosphorylated in HeLa cells in response to near-saturating EGF concentrations (7), the ratio of phosphorated EGFR to GRB2 is estimated at 1:3.5 (8). While it is clear that the absolute amounts and stoichiometry of network proteins play a major role in signal transduction (4), the impact of stoichiometric limitations on signaling dynamics is not well understood.

A recent, unexpected discovery is that RAF1, a member of the RAF family kinases that links EGFR-RAS to the mitogen-activated protein kinases (MAPKs), can be expressed at fewer than 12,000 molecules per cell (9). As a result, RAF1 abundance at the plasma membrane, which typically peaks at 10-15% of total cellular RAF1 in a very transient fashion, can be as low as 200 molecules per cell 6–8 min after EGFR activation. Yet, cells with such low-abundance RAF1 can still activate ERK robustly and with dynamics that are much more protracted than those with which RAF1 peaks at the membrane (9). Low RAF1 abundance has been observed via multiple methods in multiple cell backgrounds, including in HeLa cells (∼7800 molecules/cell) based on quantitative immunoblotting using GST-tagged standards (assuming a cell diameter of 12.9 μm) (10), and in human mammary epithelial cells (6754 molecules/cell) and triple-negative breast cancer cells (9720 molecules/cell) based on high-resolution chromatography and mass spectrometry (4). RAF1 is typically more abundant than ARAF and BRAF (4), making RAF1 the most important isoform for determining RAF’s ability to limit signal transduction.

RAS-mediated activation of RAF1 at the plasma membrane is one of the least well understood steps in the EGFR-MAPK pathway, as it is simultaneously regulated by dimerization, membrane localization, interactions with membrane phospholipids and scaffold proteins, and phosphorylation-dephosphorylation at multiple residues (11,12). The consequences of the low membranous RAF1 abundance are therefore challenging to intuit, but their identification will elucidate poorly understood aspects of pathway regulation. Low copy numbers of intracellular components generate noise in signaling pathways and cell-to-cell heterogeneity (13,14); the small number of RAF1 molecules at the membrane could play an unappreciated role in these phenomena. Low RAF1 abundance may also act as a bottleneck in ERK activation via EGFR or other RTKs and explain the poor correlation between constitutively active RAS mutants and ERK activity, which has been observed *in vitro* and *in vivo* (15–17). Finally, in settings where RAF1 is not limiting, control of ERK activity may shift from membranous RAF1 to downstream nodes.

Computational models of signaling pathways based on coupled differential equations have become a mainstay for interpreting network dynamics and for systematically identifying different operating regimes in parameter space. Parameter sensitivity analyses of the models have frequently been used to identify rate processes exerting strong control over model outputs (5,9,18–22), with some insights related to the effects of protein abundance. For example, the low abundance of Par proteins was predicted to determine sensitivity of polarized cells in *C. elegans* to chemical or mechanical perturbations (23), and low-abundance RSK was predicted and experimentally validated via immunoblot measurements to dictate the magnitude and duration of the ribosomal protein S6 kinase activity in response to erythropoietin receptor stimulation in BaF3 murine cells (24).

Many variations of parameter sensitivity analysis exist, including univariate sensitivity analysis, which quantifies changes in model output with respect to perturbations in a single parameter, and multivariate sensitivity analysis, which calculates changes in model outputs explained by simultaneous changes in multiple parameters (25,26). Sensitivity analysis is also useful in model calibration, as the analysis identifies parameters whose adjustment most impacts fit (27). This task can also be accomplished via model Sloppiness analysis, a Bayesian inference method (28) that assesses parameter uncertainty of a model constrained by experimental data and prior knowledge of parameter values (29). A byproduct of Sloppiness analysis is the identification of collective directions in parameter space that control model fit, which is particularly useful for models of biochemical networks where many parameters can vary widely and ratios of specific parameters typically control dynamics (30). Sloppiness and sensitivity analyses yield different views of parameter “importance” because the former requires experimental data (31); however, the common goal of model calibration has obfuscated this difference and led to application of the term “parameter sensitivity analysis” for both approaches (32). Key parameters identified by both methods are those whose adjustment is useful for model fitting, and parameters that are only identified by sensitivity analysis correspond to rate processes whose perturbation induces large changes in model output.

Here, we developed an ordinary differential equation (ODE)-based mechanistic model of the EGFR-ERK signaling pathway and used deterministic and stochastic solutions to gain insight into pathway operation with low RAF1 abundance. Mechanistic model analysis relied heavily on implementing and comparing a multivariate parameter sensitivity analysis with model Sloppiness analysis. The results presented, which required integration of multiple computational platforms, identified RAF1:RAS-GTP binding kinetics as critical determinants of EGFR-ERK signaling, demonstrated the ability of low-abundance RAF1 to control pathway output even with oncogenic RAS mutations, and demonstrated that stochastic RAF1 signaling dynamics arise and can propagate downstream if other nodes are also expressed at low abundance. These inferences were greatly enabled by the unique comparison of conventional sensitivity analyses with model Sloppiness analysis. The general relevance of these findings was demonstrated by an analysis of human patient tumors and cell lines showing that low RAF abundance is a common cancer cell context.

## METHODS

### Proteomic Tumor Analysis Consortium (CPTAC) Data Analysis

We used CPTAC pancreatic ductal adenocarcinoma (PDAC), glioblastoma multiforme (GBM), colon adenocarcinoma (COAD), and lung squamous cell carcinoma (LSCC) tumor proteomics data to compare relative protein expressions of EGFR, RAS (HRAS, NRAS, and KRAS), RAF (RAF1 and BRAF, where both were available), MEK1/2, and ERK1/2. CPTAC proteomics data were accessed via the *cptac* Python package (Python 3.8.5, *cptac* version 1.1.2). Percentages of tumor samples that express each protein in least amount were calculated. We used gene symbols to label the proteins encoded by each gene, as reported in CPTAC. Only samples with expression data for all proteins of interest were included in making calculations shown in Figure 1.

**Figure 1.**
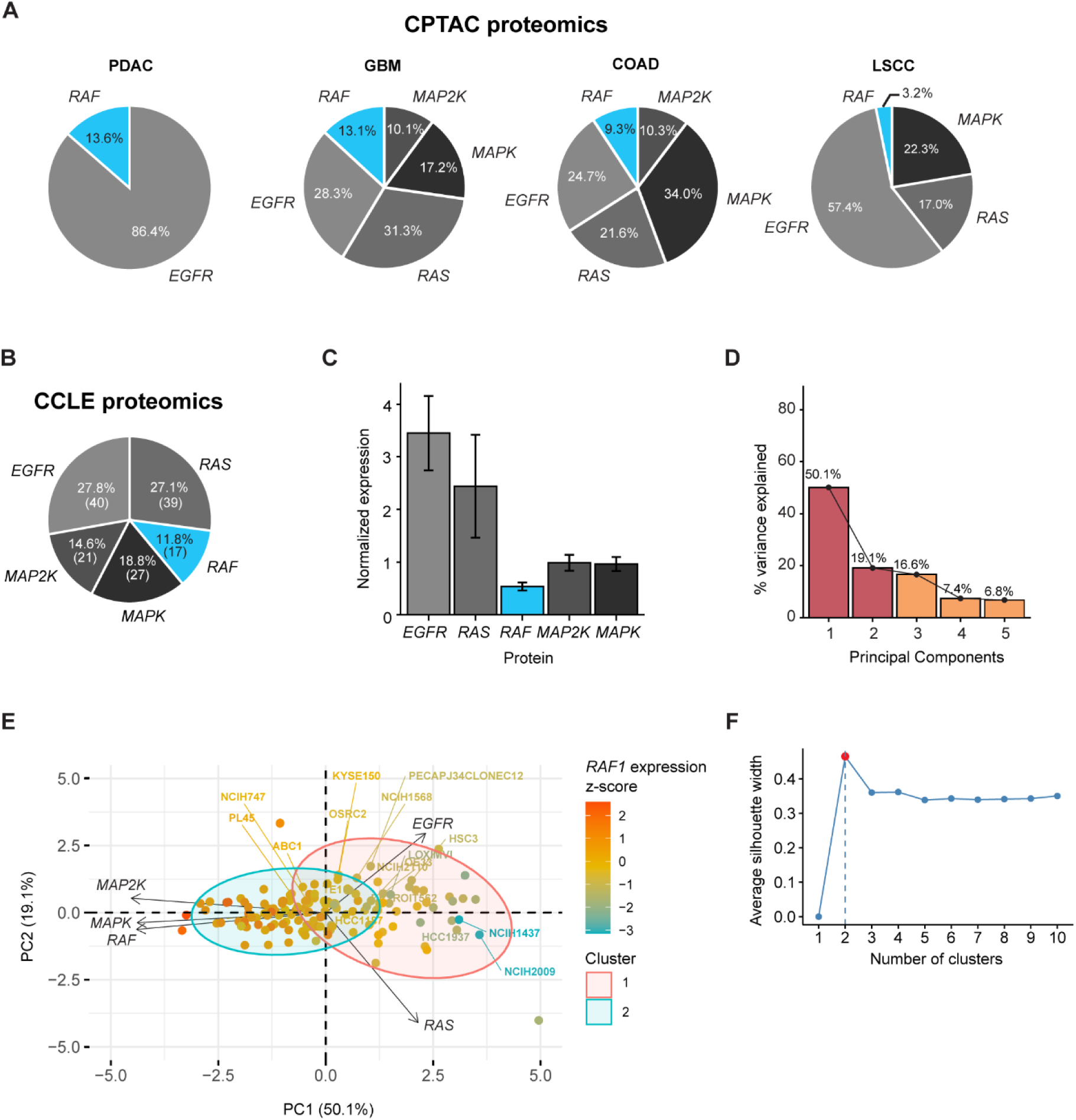
*In a substantial fraction of tumor and cancer cell settings, RAF is the least expressed EGFR-ERK pathway protein.* **(A)** Fractions of patient tumor samples from CPTAC pancreatic ductal adenocarcinoma (PDAC), glioblastoma multiforme (GBM), colon adenocarcinoma (COAD), and lung squamous cell carcinoma (LSCC) proteomics data with EGFR, RAS (HRAS, KRAS, NRAS), RAF (RAF1 and BRAF), MAP2K (MAP2K1/2), or MAPK (MAPK1/3) as the least abundantly expressed protein are shown. Proteins are labeled with the gene name that encodes for each protein, as reported in CPTAC proteomics data. **(B)** Fraction of cell lines from CCLE proteomics data that have the indicated proteins as the lowest-expressing protein is shown. Proteins are labeled with the gene name that encodes for each protein, as reported in CCLE proteomics data. **(C)** For CCLE cell lines where RAF was the least abundantly expressed protein, mean normalized expression of the indicated proteins was calculated, and standard errors of data are indicated by bars. **(D)** Mean-centered, variance-scaled EGFR, RAS, RAF, MAP2K, and MAPK expressions of cell lines from analysis in **C** were subjected to PCA. Percent variance explained by each principal component of a PCA model is shown. **(E)** For the PCA described in **D**, the PC2 vs. PC1 biplot shows the scores of each observation (cell lines, dots) colored by z-scored RAF1 expression and weights of each feature (arrows). Cell lines where RAF was the least abundantly expressed pathway protein (blue bar in **C**) are labeled. **(F)** Average silhouette scores for PCA models built with different numbers of clusters. Number of clusters used for k-means clustering followed by PCA in **E** is highlighted in red.

**Figure 2.**
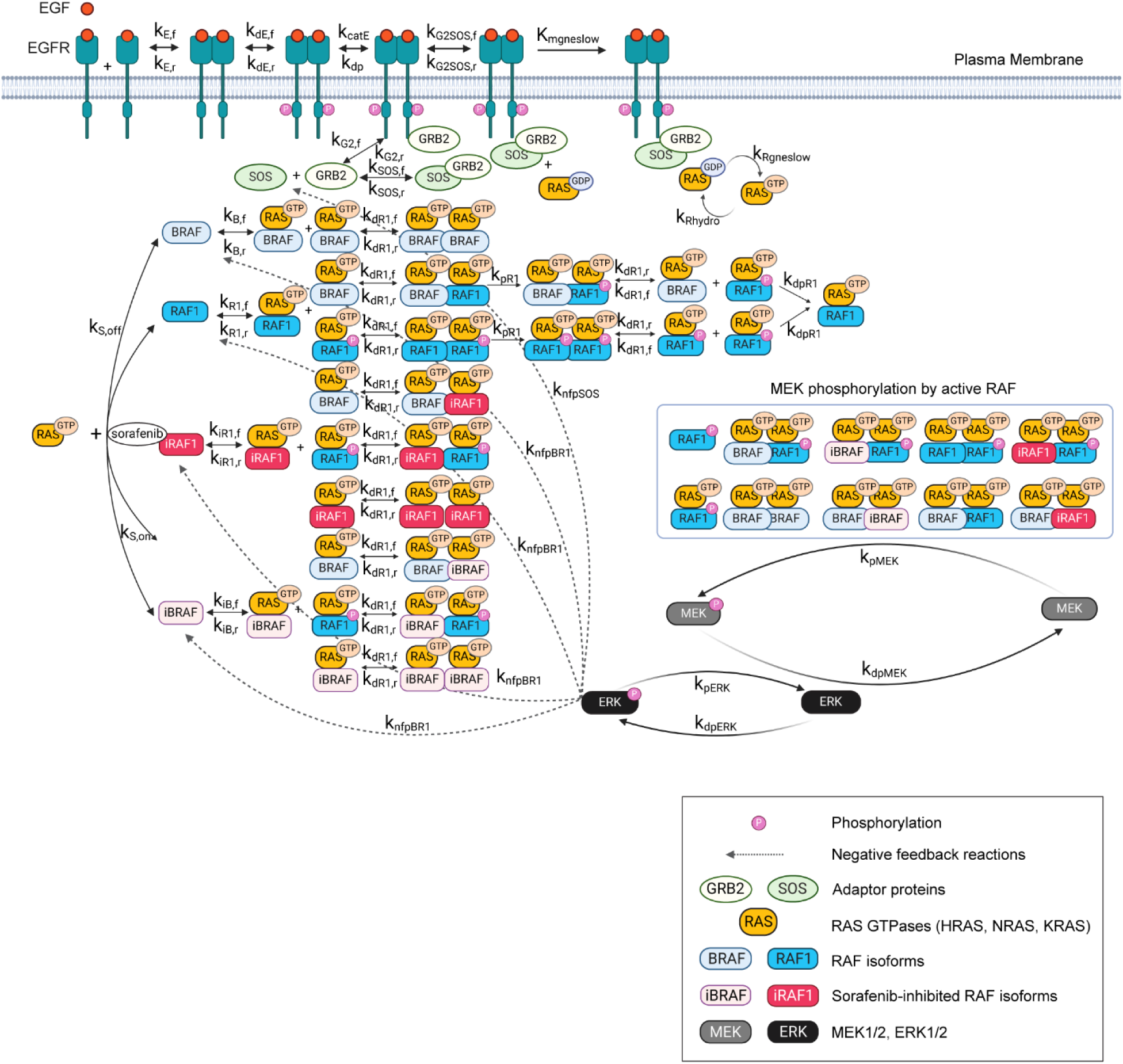
*Schematic of the EGFR-ERK signaling pathway model.* The dynamic model describes signaling initiated by epidermal growth factor (EGF) binding to its receptor, leading to EGFR dimerization and phosphorylation of a representative C-terminal tyrosine on each receptor. Phosphorylated EGFR is bound by the adaptor protein growth factor receptor-bound protein 2 (GRB2) and the guanine nucleotide exchange factor Son of Sevenless (SOS) to catalyze the exchange of guanosine diphosphate (GDP) to guanosine triphosphate (GTP) on membrane-bound RAS. GTP-loaded RAS (RAS-GTP) recruits RAF1 to the plasma membrane and induces an ‘open’ conformation for RAF1 to become phosphorylated, hetero- or homodimerize with BRAF. Catalytically active, phosphorylated RAF1 dimers recruit and phosphorylate mitogen-activated protein kinase (MEK), which catalyzes phosphorylation of extracellular signal-regulated protein kinase (ERK). Phosphorylated ERK participates in negative feedback loops, wherein ERK phosphorylates active and sorafenib-bound RAF1/BRAF (iRAF1, iBRAF), causing their release from RAS-GTP at the membrane, and SOS, disrupting its association with GRB2. Negative feedback regulations are indicated by dotted arrows. Reaction network was built in VCell, and model schematic was created with Biorender.com.

We also used the CPTAC proteomics and phosphoproteomics data to compare phosphorylated ERK levels in tumor samples of lung adenocarcinoma (LUAD) and COAD. Tumors were classified as ‘low’ or ‘high’ if RAF abundance based on expression relative to the median. Data were accessed in R programming language (version 4.2.2) via the *reticulate* package (33). Different programming languages were used because the LUAD data did not become available on CPTAC until after we performed the calculations shown here. For LUAD, RAF1 proteomics data were not available; thus, we used BRAF proteomics data. For COAD, we used RAF1 proteomics data. Proteomics data were merged with KRAS G12V mutation data to classify tumors as ‘wild-type KRAS’ or ‘mutant KRAS.’ The independent Student’s *t*-test was performed to compare the two groups based on ERK1/2 phosphorylation. Only the available phosphorylation sites in CPTAC phosphoproteomics for each tumor – ERK1 pT202/pY204 and ERK2 pT185/pY187 for LUAD, and ERK1 pT202 and ERK2 pT185 for COAD – were used for this comparison. Differences were considered significant if *P* < 0.05. All Python and R analyses are available at https://github.com/orgs/lazzaralab/repositories under the username “lazzaralab”.

See Table S1 for details on R packages and Python libraries used for analyses and Table S2 for details on the CPTAC dataset.

### Cancer Cell Line Encyclopedia (CCLE) Data Analysis

To survey RAF expression in different cancer cell settings, we used the CCLE dataset (34). Principal Component Analysis (PCA) and k-means clustering of cell lines based on CCLE mass spectrometry data were performed in R (version 4.2.2). Among 375 cell lines, 144 had complete protein expression measurements for EGFR, RAS, RAF, MEK, and ERK and were included in the analysis. Percentages of cell lines that express each protein in least amount were calculated. We used gene symbols to label the proteins encoded by each gene, as reported in the CCLE. For PCA, the optimal number of principal components (PCs) was selected as the number required to explain at least 50% of the variance in the data. k-means clustering was performed using the *cluster* package, and the optimal number of clusters to include in the PCA biplot of loadings and scores was determined by examining the number required to obtain the highest average silhouette score. *FactoMineR*, *factoextra*, and *cluster* R packages were used for figures. See Table S1 for details on R packages used for analyses and Table S2 for details on the CCLE dataset.

### Drug sensitivity and IC50 analysis

Drug sensitivity data (raw cell viability measured by Promega CellTiterGlo assay) from the Sanger Genomics of Drug Sensitivity in Cancer (GDSC)2 dataset (969 cell lines and 297 compounds) were downloaded from the GDSC project website (https://www.cancerrxgene.org/).

To assess cell sensitivity to MEK inhibitors based on RAF expression and RAS mutational status, drug sensitivity data were combined with GDSC2 genetic feature data and CCLE proteomics data and then filtered to extract results for PD0325901, refametinib, selumetinib, and trametinib (all investigational or FDA-approved MEK inhibitors). 239 cell lines with complete viability, RAF expression, and RAS mutational status data were included in the analyses. Cell lines with RAF protein expression levels below the 25^th^ percentile were categorized as “low” RAF expression; those with RAF expression above the 75^th^ percentile were categorized as “high” RAF expression. Cell lines with at least one mutant allele encoding KRAS, NRAS, or HRAS were categorized as ‘RAS mutant.’ We compared assay fluorescence intensities for cells in the presence of PD0325901, refametinib, selumetinib, or trametinib at the maximum drug concentrations tested (2.5, 10, 10, or 1 μM, respectively) that were normalized to fall within a range bounded by measurements for DMSO (control) and blank wells using the *normalizeData* function, which was included in the *gdscIC50 R* package (https://github.com/CancerRxGene/gdscIC50). The independent Student’s t-test was performed to compare the normalized fluorescence intensities of low and high RAF cell lines, with parsing based on RAS mutational status.

For analyses based on IC50 value (concentration of drug required to achieve 50% effect on viability), GDSC2 IC50 data from were downloaded from the Cancer Dependency Map (DepMap, release 24Q2). The same cell line categorizations and statistical analyses were performed to compare IC50 values for “low” and “high” RAF cell lines, again with parsing for RAS mutational status.

### Model Description

The computational model developed here used the RAS-ERK signaling dynamics model described in Surve *et al.* (9) as a foundation and describes binding interactions and reaction kinetics in the EGFR-ERK pathway. BRAF and RAF1 were the only RAF isoforms included as they are the primary binders of RAS in response to EGF stimulation (35). Based on prior absolute quantification (6), RAF1 abundance was assumed to be more than ten times greater than BRAF abundance, making it the rate-limiting RAF isoform. While the model of Surve *et al.* used a forcing function to describe RAS-GTP dynamics (9), the model developed here explicitly considered the upstream processes leading to RAS activation. This enabled prediction of how EGFR dynamics and other processes upstream of RAS affected RAF1 membrane localization in a way that was not possible with the prior model.

The model consists of 46 coupled ODEs for 46 species (Table S3) and 40 parameters (Table S4). The model describes EGF-EGFR binding, EGFR dimerization, phosphorylation-dephosphorylation of the EGFR dimer, GRB2 and SOS binding to phosphorylated EGFR, RAS activation, and negative feedback regulations of SOS and RAS-bound RAF1/BRAF molecules by ERK with or without the RAF kinase inhibitor sorafenib.

Rationale for including specific kinetic processes involved in RAF binding to RAS-GTP, RAF dimerization, phosphorylation of RAF1, MEK, and ERK, and sorafenib binding can be found in Surve *et al.* (9). Processes upstream of RAS activation were modeled as first-order, reversible reactions of EGF-EGFR binding, EGFR phosphorylation, GRB2 and GRB2-SOS complex binding to phosphorylated EGFR. RAS activation was modeled using Michaelis-Menten kinetics with RAS-GDP as the substrate and phosphorylated EGFR bound to GRB2- SOS as the enzyme that converts inactive RAS-GDP to active RAS-GTP to reflect RAS saturation by SOS and enable RAS-mediated ERK activation to be digital (36). Hydrolysis of RAS-bound GTP was modeled as a first-order, irreversible reaction. All other reactions were assumed to follow mass-action rates.

### Model Implementation

Model equations were generated using Virtual Cell (VCell) (version 7.4.0) (37,38) and were exported to MATLAB as a MATLAB-compatible ODE function, which was solved using *ode15s*, and to the SloppyCell package in Python 2 as a Systems Biology Markup Language 2 (SBML2) file (39,40). The VCell BioModel is available in the public domain at http://vcell.org/vcell-models under the shared username “LazzaraLab.” Model equations were also generated in the Julia programming language (41) (versions 1.6.2 and 1.10.0) by writing the model reactions and subsequently generating the model equations using the Catalyst.jl (42) and ModelingToolkit.jl (43) libraries, followed by solving the model equations using the DifferentialEquations.jl library (44). All MATLAB and Julia analyses are available on GitHub under the same username as above. See Table S1 for details on versions and software packages used for model analyses.

### Model Fitting in SloppyCell

The model was fit to measurements of total GTP-bound RAS with and without sorafenib, membrane-associated RAF1 with and without sorafenib, phosphorylated MEK without sorafenib, and phosphorylated ERK without sorafenib in EGF-treated HeLa cells expressing endogenous RAF1-mVenus (HeLa/RAF1-mVenus) from at least five experimental replicate measurements (64 data points total) (9,45). In the prior measurements, RAS activity was analyzed by RAS-GTP pull-down, membranous RAF1 was analyzed by live-cell imaging, and phosphorylated MEK-ERK levels were determined by immunoblot. Model fitting minimized the standard least-squares cost function, *C*(***θ***), given by

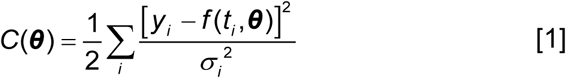

where *f*(*ti*,***θ***) represents model-predicted protein concentration for *i*th data point using the vector of best-fit parameter values (***θ***) at a time point (*ti*) and experimental time-course data point (*yi*) normalized by the variance of data point (*σi*^2^) (46).

Prior to fitting, baseline values were specified for as many rate constants as possible using previously reported values to ensure that well-known parameters did not deviate far from physiologically realistic values. Initial conditions based on literature values were fixed, and only rate constants were fitted.

Confidence intervals for SloppyCell model fits were calculated by generating a thermal ensemble of 1751 independent parameter sets to compute an average and standard deviation for model-predicted concentrations of RAS-GTP, membrane RAF1, pMEK, and pERK. For RAS-GTP, pMEK, and pERK, standard errors of the mean (SEM) based on biological replicates were available and used for generating confidence intervals. SEM for membrane RAF1 concentrations could not be quantified because membrane RAF1 was measured by quantification of live-cell images of different cells at different time points (9) In this case, Poisson errors, which scale with the number of data points, of 17% of total RAF1 concentration of 12,000 molecules were assumed for all data points.

### Model Sloppiness Analysis in SloppyCell

Stiff and sloppy parameters were identified by eigen-decomposition of the Fisher Information Matrix (FIM) as a function of changes in log-parameter values (30). Stiff parameters tend to project strongly on the FIM, meaning that changes to their values have relatively strong influence on the model’s ability to be constrained by the given data, whereas sloppy parameters have the opposite effect. However, the hallmark of a Sloppy model is log-linear eigenvalue scaling in the FIM, making it impossible to make a clear cut between “important” and “unimportant” parameters (47). To quantify model agreement with each data point achieved in each mode shown in Fig. 5G, we took the Jacobian of the cost function, J, which comprises the partial derivatives of *C*(***θ***) with respect to the log parameter values, and multiplied it with the parameter projections onto mode 0 and 1 of the FIM (46).

### Sensitivity Analyses in MATLAB

We used partial least squares regression (PLSR) as a method to conduct multivariate sensitivity analysis and refer to the approach as “PLSR SA.” While PLSR SA is not a standard multivariate sensitivity analysis approach, it is advantageous as it efficiently reduces dimensionality of feature space by identifying linear combinations of the independent variables (PLS components or latent variables) that best capture variance in related dependent variables. PLSR was performed in MATLAB using *plsregress*. For this application, the independent variable matrix was created as 3000 rows × 40 columns. Each row represented an independent set of the 40 parameters, with parameter values randomly sampled from uniform distributions spanning upper and lower boundaries created by multiplying and dividing, respectively, best-fit parameters by 10. Where indicated, the independent variable matrix was alternatively created with Latin Hypercube sampling (LHS). We used the *LHS_Call* function (26) to divide each of the parameter ranges described above into 3000 non-overlapping bins with equal probability and randomly sampled parameter values from the stratified parameter space. Once the matrix of sampled parameters was generated, entries in each column were *z*-scored to eliminate potential effects of different absolute values for each parameter. The size of the dependent variable matrix varied for different analyses. For assessing sensitivity of the EGFR-ERK system to changes in model parameters, a 3000×6 matrix was generated by aggregating the predicted maximum GTP-loaded RAS with and without sorafenib, membrane RAF1 with and without sorafenib, phosphorylated MEK, and phosphorylated ERK concentrations in response to 10 ng/mL EGF for each parameter set. For evaluating sensitivity of ERK activity in mutant RAS relative to wild-type RAS to changes in model parameters, a 3000×1 matrix was generated by computing the difference in time-integrated phosphorylated ERK concentrations between wild-type and mutant RAS for each parameter set. Entries in each column of the dependent variable matrices were z-scored. A six-component PLSR model was chosen to maximize fit (R^2^X, R^2^Y) and predictive power (Q^2^Y) with the number of components matching the number of modes in Sloppiness analysis.

For “filtering” parameter sets to achieve better agreement between PLSR SA and Sloppiness analysis shown in Fig. S1A-E, parameter sets used in PLSR SA were ranked in descending order by the cost *C*(***θ***). The top third of the parameter sets with smallest aggregate error were retained, and PLSR was performed using the six time-integrated model outputs described above. Here, the dependent variable was changed from maximum to time-integrated model outputs because the model cost basin required for Sloppiness analysis is defined more closely by time-integrated model output than the maximum output.

Univariate SA (i.e., parameter perturbation one-at-a-time), was conducted by individually increasing and decreasing parameter values by a factor of 10 from their best-fit values. Sensitivity was calculated by summing the integrated differences between the model output generated by baseline parameter values and that by perturbed parameter values.

### Model Fitting/Parameter Estimation in MATLAB

We examined efficiency of model fitting as a function of different numbers of parameters varied in MATLAB. Model fits obtained by varying all versus a subset of important parameters nominated by PLSR SA were compared. From both fitting scenarios, we obtained a single set of best-fit model predictions. For selecting a subset of parameters, PLSR SA was performed using the independent variable matrix described in *Sensitivity Analyses in MATLAB* and the cost function values as the dependent variable. Parameters with variable importance in projection (VIP) scores > 1 were retained for model fitting. From fitting only the VIP > 1 parameters, we additionally obtained multiple model predictions with confidence intervals. The same convergence criterion was used for the two fitting scenarios such that the *fminsearchbnd* constrained Nelder-Mead optimization routine continued until the change in cost between iterations was < 1×10^-4^, with a maximum allowable number of iterations of 4×10^4^. Signaling dynamics predicted from the best fits obtained in SloppyCell and MATLAB are shown in Figures 3 and 5D, respectively. Performances of the two methods were also compared in Table S5 in terms of costs for the single best fits.

**Figure 3.**
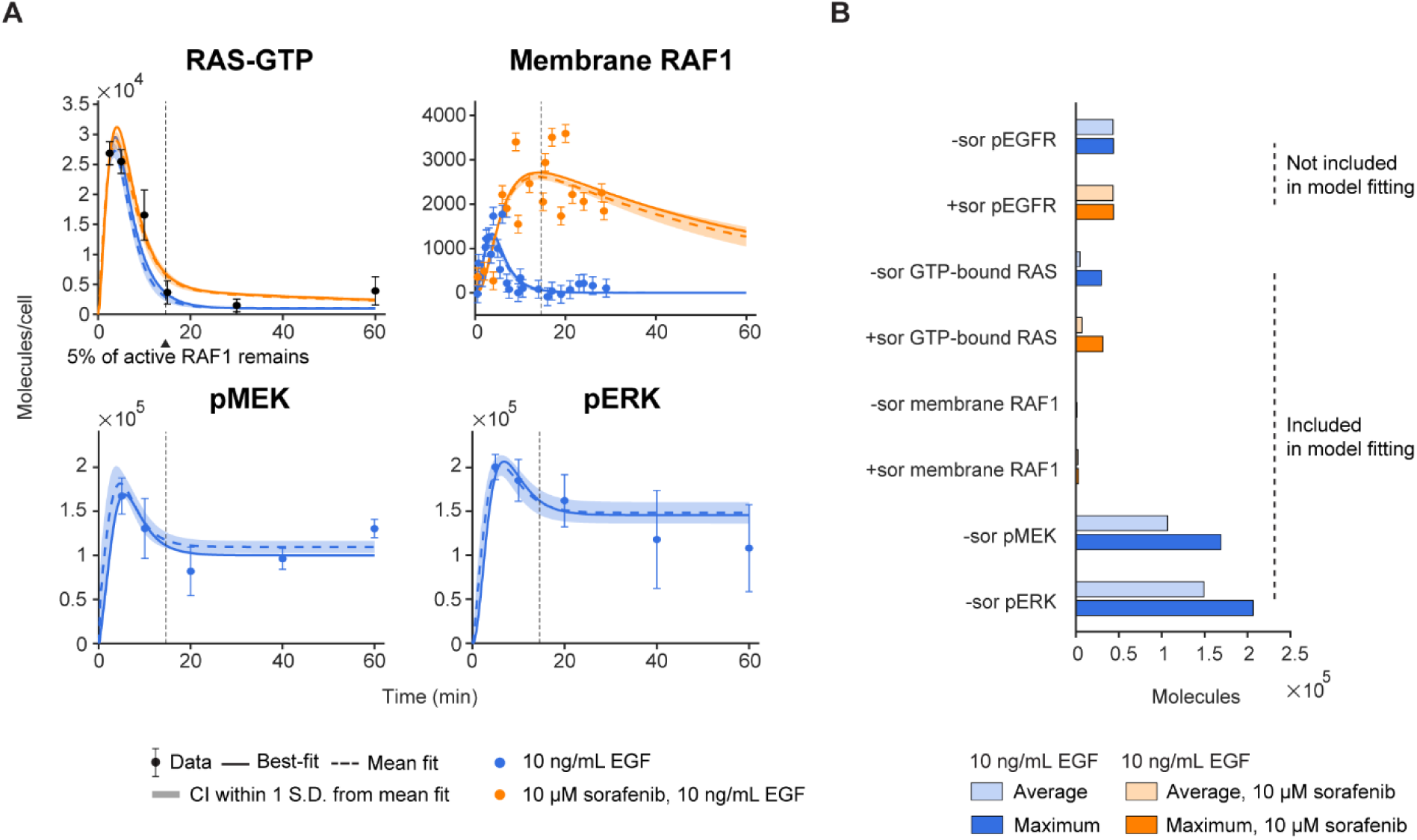
*Fitted model recapitulates signaling dynamics measured in EGF-treated HeLa cells*. **(A)** Model rate constants were fitted against immunoblot measurements for RAS-GTP in HeLa/HRAS-mVenus cells reported by Pinilla-Macua *et al*. (45) and live-cell imaging of membrane RAF1 with or without 10 μM sorafenib and immunoblot measurements for pMEK and pERK in HeLa/RAF1-mVenus reported by Surve *et al*. (9). Time points at which 5% of RAF1 remained active at the plasma membrane are indicated by vertical dotted lines. Throughout the panels, the means of model outputs generated from 1751 parameter sets are indicated by dotted lines, best-fit model outputs are indicated by bold lines, 68% confidence intervals (CI) about the means are represented by shaded areas, data are indicated by colored dots, and standard errors of data are indicated by bars. **(B)** Predicted maximum and time-averaged concentrations of the indicated proteins in response to 10 ng/mL EGF for 60 min with or without 10 μM sorafenib from the best-fit model. pEGFR includes pEGFR, pEGFR bound to GRB2, and pEGFR bound to GRB2-SOS, RAS-GTP includes all RAS-GTP bound species, membrane RAF1 indicates all RAF1-bound species at the plasma membrane. Vertical cross bars indicate whether the dataset was included in model fitting. Model fitting was performed in SloppyCell.

### Confidence Interval Calculations

Where shown, confidence intervals for model fits were generated using parameters nominated by PLSR SA as strong controllers of model output. This substantially reduced the computational time required. Generation of data sets was performed by Monte Carlo simulation in MATLAB by assuming a normal distribution of each experimental data point with the mean and standard deviation at each time point as reported in the experimental data used for fitting (9,45). For each time point, 20 data points were generated by random sampling. The data points were used in minimizing the cost function (Eq. [1]) by fitting the model 20 times with the *fminsearchbnd* in MATLAB. The collection of fits was used to compute 68% confidence intervals of with degrees of freedom of 99 based on two-tailed student’s *t* distribution, using the MATLAB Statistics Toolbox *tinv* function.

### Comparison of PLSR SA and Sloppiness Analysis

PLSR SA and Sloppiness analysis results were compared by scaling PLSR SA parameter loadings to match the log range of projections onto each mode in the FIM (-0.6∼0.6) from Sloppiness analysis. The number of PLS components and modes to compare was determined by finding the total number of PLS components that produced a cumulative predictive power > 50% and the number of stiff modes at which eigenvalues of the FIM (λ) were > 0. For comparing aggregate importance of parameters across PLS components in PLSR SA and modes in Sloppiness analysis, we first calculated VIP scores of PLSR SA for up to six components. The equivalent metric in Sloppiness analysis was calculated by taking the squares of individual parameter’s projection in modes up to 5, with projection in each mode weighted by the corresponding λ. Similarity between the magnitudes of projection of each parameter onto each mode in Sloppiness analysis and PLS component in PLSR SA was calculated using the *corr* function in MATLAB, which provides Pearson’s correlation coefficients and their associated *P*-values using two-tailed Student’s *t*-test. Similarity between the two methods was also calculated using the Jaccard index, defined as

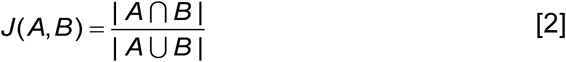

where *A* and *B* represent a set of parameters that have scaled sensitivity values above a threshold value in each PLS component in PLSR SA and mode in Sloppiness Analysis.

### Stochasticity Sensitivity Analysis in VCell and Julia

Sensitivity analysis of stochastic simulations was performed by translating the VCell EGFR-ERK BioModel to a VCell MathModel, followed by importing a LHS parameter array with 198 total parameter sets (upper limit of the number of stochastic simulations/parameter batch array in VCell at the time the simulations were performed) using the Batch Simulation feature (available in v7.6.0), with parameter upper and lower bounds set as described above for PLSR SA parameter sets. Five stochastic simulations were then performed for each LHS-generated parameter set (N=5) in VCell using the Gibson Next Reaction Stochastic solver, and the stochastic model outputs were exported to Julia for sensitivity analysis. PLSR SA of the VCell- generated stochastic outputs of maximum membrane RAF1 level was performed in Julia using ScikitLearn.jl (https://github.com/cstjean/ScikitLearn.jl). Confidence intervals for the PLSR coefficient estimates were generated via the bootstrap method, wherein the observations (198×40 matrix of randomly sampled parameter values and the model outputs corresponding to each parameter set) were drawn 10,000 times with a resampling fraction of 0.9. The MathModel with 198 LHS parameter arrays is available in the public domain at http://vcell.org/vcell-models under the shared username “LazzaraLab.”

We noticed that having too few parameter sets often resulted in variation in predictive power among multiple PLSR SA runs, causing PLSR SA results to be “unstable.” To ensure stability of the stochastic PLSR SA results, we increased the number of LHS-generated parameter arrays to 1000, which consequently required 1000 stochastic simulations and an inordinate amount of simulation time on the VCell’s shared server. Thus, we recreated the EGFR-ERK model in Julia as described above and generated model solutions using the Sorting Direct Method (48). Five stochastic simulations for each of the 1000 parameter arrays were distributed across 40 cores on the UVA High Performance Computing cluster, Rivanna, significantly reducing the computational cost of model simulations compared to VCell’s batch simulations (∼0.08 min per single trajectory stochastic run on Rivanna vs. >1 min on VCell). The model predictions from each set of five simulations were averaged, and a 3000×1 matrix of membrane RAF1 stochasticity scores, defined in *Results*, for each parameter set was used as the dependent variable. Confidence intervals for the PLSR SA VIP scores and coefficient estimates were generated via the bootstrap method described above.

### Other Sensitivity Analyses in Julia

Other multivariate sensitivity analyses—specifically the LHS-partial rank correlation coefficient (PRCC) (26) and extended Fourier Transform Sensitivity Test (eFAST) methods (49)—and derivative-based local sensitivity analysis were performed in Julia using the GlobalSensitivity.jl (50) and SciMLSensitivity.jl (https://github.com/SciML/SciMLSensitivity.jl) libraries, respectively. For eFAST, changes in maximum and average model outputs for each pathway node considered in model training were calculated as functions of simultaneous changes in all rate constants within an order of magnitude above and below the baseline values. For each parameter, we calculated first-order eFAST indices, defined as variance in model output due to that particular parameter of interest, and total-order eFAST indices, defined as the sum of first-order index and variance in model output due to nonlinear interaction between that parameter and other parameters (26).

## RESULTS

### Low RAF abundance is characteristic of a substantial fraction of tumors and cancer cells

To assess the frequency of low RAF abundance in settings of interest, we examined relative expression of the key EGFR-ERK pathway proteins using CPTAC mass spectrometry data. Rank-ordering the expression of EGFR, RAS (HRAS, KRAS, NRAS), RAF (RAF1 and BRAF), MEK1/2, and ERK1/2 across tumors of different types revealed that 14% of PDAC, 13% of GBM, 9% of COAD, and 3% of LSCC patients expressed RAF as the least abundant protein (Fig. 1A). Performing the same analysis using CCLE mass spectrometry data, 11% of cell lines with complete expression profiles of the queried proteins had RAF as the least abundant protein, 4.5-fold lower on average than the other proteins assessed (Fig. 1B,C). A PCA projection of the CCLE cell lines with EGFR, RAS, RAF, MEK, and ERK as features fascinatingly demonstrated that cells separated along an axis spanning two overall groups – cells with relatively high expression of MAPK cascade proteins (RAF, MEK, ERK) and low expression of EGFR and RAS or those with relatively high expression of EGFR and/or RAS but low MAPK cascade protein expression (Fig. 1D-F). Based on that observation, performing k-means clustering with k = 2 revealed a cluster substantially enriched for cell lines where RAF proteins were the least abundant. Thus, low RAF abundance is a defining characteristic of a substantial fraction of tumors and cancer cell lines and may be expected to play the role of a stoichiometric bottleneck in those settings. When RAF abundance is low, it is particularly low relative to EGFR and RAS upstream. That fact and its position as the initiating kinase in the three-tiered MAPK cascade make RAF unique as a low-abundance protein.

### The fitted network model captures RAF-limited EGFR-ERK pathway dynamics

To understand the impact of kinetic perturbations on EGFR-ERK signaling dynamics in the context of low RAF1 abundance, we constructed a computational model that describes activation of the EGFR-ERK pathway using coupled ODEs. A schematic showing model species (Table S3) and interactions among them is presented in Fig. 2. Rate terms in the model equations were based primarily on mass-action considerations used to model the continuum behavior of these types of systems. We first fit our model to previously reported data from HeLa cells with fluorescently tagged endogenous HRAS or RAF1 for total GTP-bound RAS, membrane-associated RAF1, and whole-cell pMEK1/2 and pERK1/2 in the presence or absence of the RAF inhibitor, sorafenib (9,45) (Fig. 3A). The availability of measurements with sorafenib strengthens model training by providing a condition with a RAF perturbation. When sorafenib was absent, GTP-bound RAS and membranous RAF1 peaked 2.5 to 5 min after EGF treatment, respectively, while pMEK and pERK displayed sustained activation at least through the first hour of response. Standard deviations of pMEK and pERK were relatively large, and membrane-associated RAF1 at later times lacked any training data, resulting in large confidence intervals compared to earlier time points. Unlike the activities measured by immunoblot at other pathway nodes, membrane RAF1 levels were measured by live-cell microscopy using smaller time intervals (0.25 - 3 min) for less overall time (30 min after EGF addition) to prevent photobleaching. The tendency for RAF1 to remain associated with the membrane for longer times in the presence of sorafenib, which can be seen in the second panel of Fig. 3A, has been documented and explained (9,51,52). Based on that prior work and because sorafenib is a specific inhibitor of RAF, but not RAS, we assumed that GTP-RAS loading was unchanged by sorafenib and therefore used the same data points for GTP-RAS level with and without sorafenib for model training.

As described in Methods, we attempted to fit the model using two separate approaches. We used the best-fit parameter values from SloppyCell, obtained by a combination of Nelder-Mead derivative-free (53) and Levenberg-Marquardt derivative-based (54,55) optimization, because it produced a smaller cost function value than a fit we performed in MATLAB using only Nelder-Mead (Table S5). Quality of model fit was reflected by the good order-of-magnitude comparison between the SloppyCell-predicted confidence interval bounds and the experimental errors (average difference of 54%; Fig. 3A). Note that the small model-predicted differences in GTP-RAS levels with or without sorafenib result from minor differences in the fitted values of the four parameters that pertain to the presence of sorafenib (*kiBf*, *kiBr*, *kiR1f*, *kiR1r*) compared to their respective fitted values in the absence of sorafenib (Table S4). Comparison of the model-predicted maximum and time-averaged concentrations of the six signaling species used in model training highlights the low abundance of RAF1 at the membrane compared to other species and a relatively small difference between maximum and time-averaged levels of pMEK and pERK species, which exhibit protracted activation (Fig. 3B).

### Low membrane RAF1 abundance limits EGFR-ERK pathway flux

To identify the key determinants of EGFR-ERK signal transduction in the low-RAF1 context, we performed model Sloppiness analysis and multivariate sensitivity analyses using partial least squares regression (referred to as PLSR SA) with dependent variables set as the six pathway nodes used for parameter estimation (Fig. 4A). PLSR SA is described in *Methods* and was used here as a type of multivariate sensitivity analysis to identify the parameters that most strongly controlled outputs of interest without constraint on how much the output varied. In contrast, Sloppiness analysis determines sensitivity of the model fit to experimental data for changes in parameter values. A direct comparison between multivariate sensitivity analysis and Sloppiness analysis for the same model and data has not previously been performed, to our knowledge, but it is useful because parameters that control a model fit versus robustness of output to parameter perturbations may differ in non-intuitive ways. We first compared parameter loadings in PLSR SA scaled to the range of parameter projections on the FIM in Sloppiness analysis (-0.6∼0.6) and defined the interval as *scaled sensitivity*. This revealed some general similarities and differences between sensitivity and Sloppiness analyses (Fig. 4A). Many, but not all, of the parameters that exhibited large magnitudes of scaled sensitivity, and VIP scores in PLSR SA, which is the sum of each parameter’s loadings across all PLS components weighted by the change in model outputs explained by that parameter, were also nominated as important parameters by Sloppiness analysis. In some cases, the directionality of the computed effects was different though between the two methods. The major difference was that the matrix of scaled sensitivities in Sloppiness analysis was much more sparsely populated by values that deviated substantially from ∼0, presumably because Sloppiness intrinsically accounts for the ability of other parameter variations to compensate for changes in one parameter and still achieve a reasonable fit.

**Figure 4.**
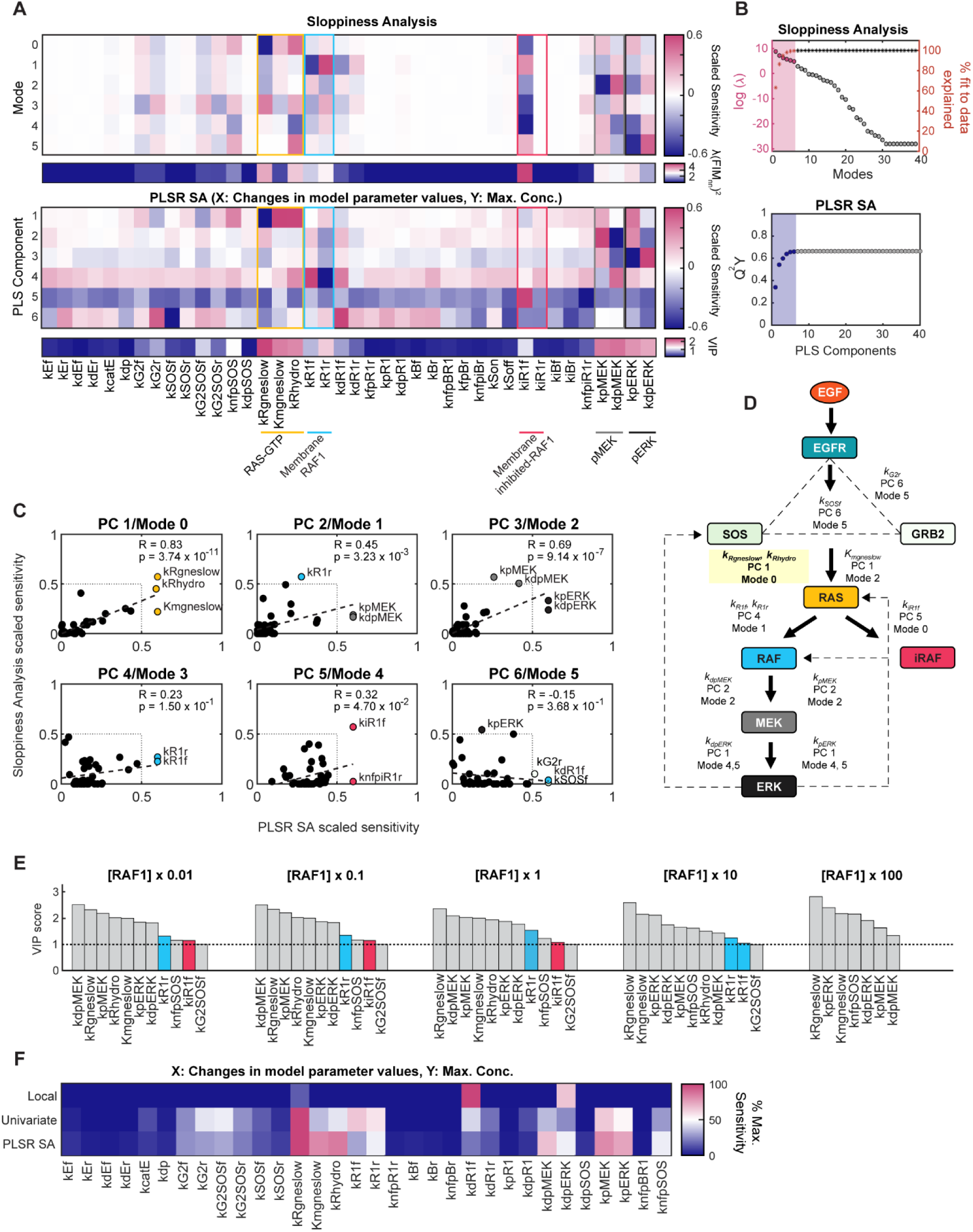
*EGFR-ERK signaling system behavior is primarily controlled by RAS activation and RAF1:RAS-GTP unbinding rates.* **(A)** Sloppiness analysis was conducted by eigen- decomposition of the FIM as a function of log-parameter values from the 1751 parameter sets described in Fig. 3. PLSR SA was performed using 3000 uniformly sampled, random parameter sets (3000 rows × 40 columns) as the independent variable matrix and predicted maximum concentrations of RAS-GTP with or without sorafenib, membrane RAF1 with or without sorafenib, pMEK, and pERK in response to 10 ng/mL EGF from each parameter set (3000 rows × 6 columns) as the dependent variable matrix. Projection of parameters onto the first six modes from model Sloppiness analysis (top) and parameter loadings in the first six PLS components from PLSR SA (bottom) were compared. PLSR SA parameter loadings were scaled to the range of projections of the FIM in Sloppiness analysis (-0.6∼0.6) and plotted as scaled sensitivity. For each parameter, aggregate weights across modes in Sloppiness analysis (λ(FIMnn)^2^), calculated by adding the squares of projection onto each mode (FIMnn) weighted by the corresponding eigenvalues (λ), and VIP scores from PLSR SA were also plotted. **(B)** Natural log of eigenvalues (λ) of the FIM matrix, which indicates sensitivity of cost *C*(***θ***) to perturbations along orthogonal directions in parameter space, and percent model fit explained by each mode from Sloppiness analysis for increasing numbers of modes (top) and cumulative predictive powers (Q^2^Y) of PLSR SA model for increasing numbers of PLS components. Selected numbers of modes and PLS components for comparison are highlighted (bottom). **(C)** Scatter plots of scaled sensitivities in Sloppiness analysis and PLSR SA are shown. Pearson’s correlation coefficients (R) of scaled sensitivities are calculated for each respective component and mode. Parameters with scaled sensitivity from both analyses > 0.5 are labeled and filled with color. *P* values were calculated using one-tailed Student’s *t*-test. **(D)** Qualitative summary of panels **A** and **C**. Parameters that exerted relatively strong control over the maximum concentrations of the six model outputs are depicted on a map of key pathway nodes of the EGFR-ERK network. The parameters identified to have strongest control over the EGFR-ERK signaling system by both PLSR SA and Sloppiness Analysis are bolded and highlighted in yellow box. Protein-protein interactions are shown as dotted lines, inhibition of RAF1 by sorafenib is shown as a “T” line. **(E)** PLSR SA was performed using the same independent and dependent variables as in **A** with indicated perturbations to baseline RAF1 abundance. Rank- ordered VIP scores of model parameters from PLSR SA for each RAF1 abundance were calculated. Parameters with VIP scores > 1 are shown. **(F)** Comparison of local, univariate, and PLSR SA with dependent variables set to predicted maximum concentrations of GTP-RAS, membrane RAF1, pMEK, and pERK in response to 10 ng/mL EGF. Changes in the model outputs, normalized by the maximum change for each analysis, were calculated.

**Figure 5.**
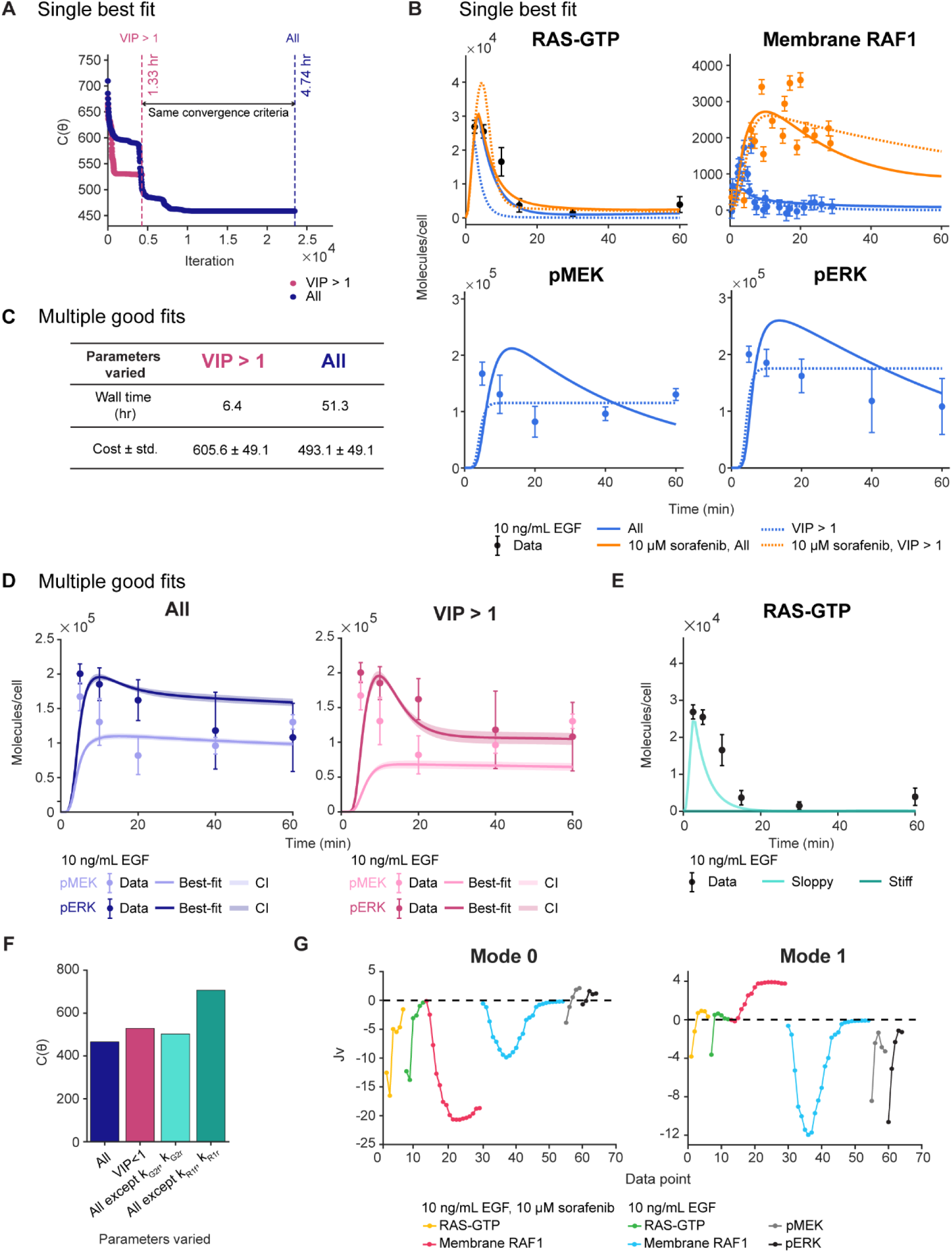
*RAF1:RAS-GTP binding parameters control model accuracy and efficiency of model fitting*. **(A)** The model was fitted by allowing all 40 parameters or a subset of parameters with VIP scores > 1 from PLSR SA in Fig. 4A to vary. Convergence of cost function value *C*(***θ***) over iterations and corresponding computational wall time (total time required for the optimization to run) are shown. The vertical dashed lines show the position of iterations corresponding to the wall times required to satisfy the convergence criteria described in *Methods*. **(B)** Model fit results from the two cases described in **A**. In the model fitted with a subset of parameters varied (VIP > 1), 11 parameters of importance nominated by PLSR SA in Fig. 4A (*k_pMEK_, k_nfpSOS_, k_dpERK_, k_R1r_, kRgneslow, kG2SOSf, kiR1f, kpERK, kRhydro, Kmgneslow, kdpMEK*) were varied. **(C)** Summary of computational time and minimum cost reached for varying all parameters versus only the subset of parameters when model was fitted to a synthetic dataset that consisting of 20 data points generated based on a normal distribution about the mean. In the latter case, the same subset of parameters with VIP > 1 as in **A** were allowed to vary. **(D)** Representative pMEK and pERK model fits from the two cases described in **C**. Shaded areas indicate 68% confidence intervals obtained by assuming Gaussian distribution of data about the means. **(E)** Representative RAS-GTP fits when the stiff parameters, RAF1 activation parameters (*kR1f, kR1r*), identified from Fig. 4A were fixed to base values while the other parameters were varied, or the sloppy parameters pTyr- SH2 domain interaction parameters (*kG2f, kG2r*) were fixed to base values while the other parameters were varied are shown. **(F)** Minimum values of the cost function *C*(***θ***) are compared between model fits with indicated parameters varied. **(G)** Products of the Jacobian (J), which is the partial derivatives of the cost function *C*(***θ***) with respect to the log parameter values, and the parameter projection onto mode 0 and 1 (v) from Sloppiness analysis in Fig. 4A, indicated as Jv, against individual data points for indicated key pathway nodes included in model fitting from Fig. 3A are shown.

We surveyed the first six PLS components (sometimes referred to as latent variables) from PLSR SA and the first six modes from SloppyCell because these components and modes explained the majority of variance in model outputs in PLSR SA (>50%) and in model fit to data in Sloppiness analysis (>99%), respectively (Fig. 4B). Log-linear scaling of the eigenvalues λ of the FIM over four orders of magnitude indicated that the EGFR-ERK model system was indeed Sloppy (Fig. 4B). We then compared parameter loadings in PLS components and Sloppiness modes because both inform the relationship between parameters and model behaviors — as linear combinations of parameters that control model outputs for PLSR SA, and as products of parameters (linear combinations of log parameters) that control the FIM for Sloppiness analysis (56). Correlation between scaled sensitivities of parameters that overlapped in the two analyses was the strongest for the first component/mode (Fig. 4C). RAS-related parameters (*k_Rgneslow_*, *kRhydro*) were identified as the most critical regulators of the EGFR-ERK system by the first (most important) PLS component/mode. RAF1:RAS-GTP (un)binding parameters (*kR1f*, *kR1r*) were heavily weighted in the second mode, and MEK (de)phosphorylation parameters (*kpMEK*, *kdpMEK*) were heavily weighted in the second PLS component.

In search of methods that would lead PLSR SA to behave more like Sloppiness analysis, we repeated the comparison using only the 1000 random parameter sets for PLSR SA that yielded the lowest rank-ordered cost values (Fig. S1A). The importance of RAF1:RAS-GTP binding parameters was accentuated in PLSR SA (Fig. S1B). We selected 1000 parameter sets, as fewer parameter sets failed to produce stable PLSR loadings and predictive power (Fig. S1C,D). With this cost-value filtering approach, the overlap between influential parameters identified by PLSR SA and Sloppiness analysis was improved, indicated by a larger Jaccard index of unique parameters that have scaled sensitivity values above 0.3 (Fig. S1E). The strength of the linear relationship between each PLS component and mode was weaker in the filtered parameter case, potentially because the lower-ranked PLS Components had more weight in explaining the model fit than the variance in model outputs and the components are not altered to be linear combinations of products of parameters as in Sloppiness modes by the filtering approach (Fig. S1F). Overall, this comparison reveals that RAS- and RAF1-related parameters control EGFR-ERK signaling dynamics within and outside the range of the data observations used for model fitting.

We further noted that dysregulation of the EGFR-ERK system, modeled by large changes in parameter values in PLSR SA, shifted the control of the signaling dynamics to rate processes downstream of RAF1. We observed this shift in the second-most important PLS component/Sloppiness mode, where Sloppiness analysis emphasized the importance of RAF1:RAS-GTP unbinding parameter (*kR1r*) while PLSR SA highlighted the importance of MEK (de)phosphorylation parameters (*kpMEK, kdpMEK*) in controlling the dynamics of the six core proteins in the pathway (Fig. 4C). A qualitative summary of the PLS components and modes onto which key parameters in the reaction network most strongly project shows that, when the range of parameter variations is not restricted by how well the model fits to data (as in PLSR SA), control of the EGFR-ERK system behavior shifts from RAF to MEK and ERK (Fig. 4D). Note that numbering of modes and PLS components begins at 0 and 1, respectively. Thus, the PLS component numbers match the corresponding mode numbers for upstream (EGFR to RAF) levels of the pathway, but the correspondence between PLS components and Sloppiness modes decreases further down the cascade.

To understand the degree to which low RAF1 abundance may present potential stoichiometric limitations in EGFR-ERK signal transduction, we calculated VIP scores from PLSR SA of the fitted model for different RAF1 expression levels (Fig. 4E). Interestingly, RAF1:RAS-GTP binding parameters were no longer of importance when RAF1 abundance was increased 100-fold, indicating that EGFR-ERK system dependence on membrane RAF1 levels is characteristic of the low-RAF1 abundance setting. Local sensitivity analysis (partial derivatives of an output with respect to the small changes in a parameter of interest (57)) revealed that the overall signaling dynamics was only sensitive to small changes in RAF dimerization (*kdR1f*) and ERK dephosphorylation rates (*kdpERK*) around each of their fitted values, and univariate sensitivity analysis, which quantifies the effects of a fixed range of perturbation in individual parameters on the combined model outputs, de-emphasized the importance of MEK and ERK dephosphorylation rates (*kdpMEK*, *kdpERK*) compared to PLSR SA (Fig. 4F). This suggests that the overall EGFR-ERK signaling dynamics are more sensitive to large, step changes in RAS and RAF1-related rates or simultaneous changes with other rates than those in downstream parameters.

### RAF1 activation parameters control model fit and filter out upstream system perturbations

The biological insights gained from PLSR SA can also be used to improve the efficiency of model fitting, but with tradeoffs in fit quality. We demonstrated this by comparing model fits in MATLAB permitting simultaneous variations in all 40 kinetic rate constants or the subset of rate constants with PLSR SA VIP scores > 1 identified in Fig. 4A (Fig. 5A). Model fit by varying only high-VIP parameters, including those describing RAS activation, ERK (de)phosphorylation, and RAF1:RAS-GTP (un)binding, yielded a prediction error that was ∼25% larger than that achieved using all parameters but with a 75% reduction in computational time when the same convergence criterion was applied (Fig. 5A,B, Table S6). We tested LHS, a pseudo-random method that is generally preferred over random sampling in model fitting (26), to select parameters to vary in model fitting but did not see a clear benefit. This surprising result may have arisen due to the large number of parameter sets needed for a stable PLSR model (i.e., without variation in predictive power among multiple simulation runs) overwhelming any potential benefit of alternative sampling approaches (Table S7, Fig. S1G). The difference between the two model fits in Fig. 5A, B likely stems from the reduced degrees of freedom in the fit when only a subset of parameters was allowed to vary. The efficiency in fitting gained by parameter filtering becomes especially apparent when obtaining distributions of fits, rather than a single best-fit result. Indeed, as shown in Fig. 5C, D, an 8-fold improvement in wall time with only a 23% increase in the cost function was achieved with parameter filtering based on PLSR SA when we sought the 20 best fits that accounted for the experimental error in the model training data. Results are shown in Fig. 5C for pMEK and pERK only because these results best exemplify the extremes in alternative fit results.

In addition to issues of computational cost, there are other reasons why the choice of varying all or a subset of model parameters should be made with caution. Model fitting in SloppyCell was performed by varying all parameters simultaneously for fixed initial conditions, including the RAF abundances, based on the premise that most parameters are unknown and that obtaining precise estimates is less important than achieving high accuracy of predictions (47,58). However, cost function minimization achieved by varying all parameters typically came at a cost of some fitted parameter values being incongruous with known values (Fig. S2). For example, the implied equilibrium dissociation constant for pEGFR binding to GRB2 of 0.002 µM from the SloppyCell fit was inconsistent with the experimentally determined range of ∼0.03∼0.06 µM (59). This and other such discrepancies are shown in Table S7. Importantly, Sloppiness analysis can aid in determining whether such deviation matters. Fixing Sloppy parameters (i.e., those with weak projection onto modes 0 or 1), such as those for pTyr-SH2 interaction (*kG2f*, *kG2r*) (Fig. 4A) to biologically known values had negligible effect on model fit when the remaining parameters were allowed to vary. Conversely, moving the stiff RAF1:RAS-GTP (un)binding parameters (*kR1f*, *kR1r*) to model-fitted values reported previously (60) led to an obviously erroneous fit result that produced virtually no RAS activation (Fig. 5E,F). This was consistent with the importance of moving RAF1:RAS-GTP (un)binding parameters in directions specified by Mode 1 to membrane RAF1 fit to data, revealed by Sloppiness analysis (Fig. 5G).

To understand how perturbations in rate processes control signaling at four levels of the cascade, from RAS to ERK, we compared different types of multivariate SA to univariate and local SA against individual model outputs. Univariate SA revealed that the maximum level of active species at each tier of the cascade was only controlled by the nearby activation rates of the species (*kRgneslow*, *kR1f*, *kpMEK*, *kpERK*). For example, ERK activity was controlled only by ERK- proximal processes. Conversely, multivariate PLSR SA demonstrated that control of active RAS and RAF1 levels was distributed throughout the signaling cascade (Fig. S3A). Membrane RAF1 level was especially sensitive to rate processes governing all four levels of the cascade (*kR1r, kRhydro, knfpSOS, kR1f, kpERK, Kmgneslow, kpMEK*), highlighting the role of membrane RAF1 as the most critical determinant of system performance. A similar distribution of control over GTP-RAS and membrane RAF1 was observed in sensitivity analysis using a PRCC approach (Fig. S3B). Using a local SA approach, we observed that the rate constant for ERK phosphorylation exerted the strongest local control over ERK activity (Fig. S3C), consistent with previous findings that local changes in ERK phosphorylation rates and degree of multisite phosphorylation (i.e. processivity) by mutations in MEK can induce distinct ERK activation dynamics and phenotype (61). As a final way to investigate system sensitivity, eFAST indices were computed (Fig. S3D,E) (62). Total-order eFAST indices were generally much larger than first-order indices, suggesting the importance of nonlinear interactions among rate processes in system regulation. Moreover, when interactions among parameters were considered through total-order calculations, RAS- and RAF1-related parameters were again among the most important.

### Low membrane RAF1 expression dampens ERK activity driven by oncogenic KRAS

The emphasis of parameter sensitivity results on RAS- and RAF1-related parameters led us to question if RAF1 acts as a bottleneck when RAS is mutated. Constitutively active RAS mutants are generally thought to drive cancer at least in part by increasing RAF/MEK/ERK signaling, but *in vitro* and *in vivo* studies suggest that ERK activity is not always well correlated with RAS mutation (16,17). KRAS is the most abundant RAS isoform in many cell backgrounds and the second most abundant in HeLa cells (after HRAS) (4,6). Thus, inferences from a model based on the expression of all RAS isoforms may be reasonably expected to reflect scenarios where KRAS is mutated. KRAS mutation is the most common genetic abnormality in cancer, with G12D, G12V, G13D, and G12C mutants accounting for 83% of all clinically observed mutants (63). We focused on KRAS^G12V^ (the second-most common KRAS mutation in colorectal, lung, and pancreas cancers) because it has the largest difference in hydrolysis rate constant compared to wild type among the above four most common KRAS mutations, which is advantageous in predicting an upper limit for KRAS mutant effects (64,65). To model KRAS^G12V^ expression, the rate constant for RAS-GTP hydrolysis was reduced 16-fold to match the previously reported GTP hydrolysis rate for the KRAS^G12V^ mutant (64). No retraining of the model was performed.

The model predicted only a ∼1.3-fold difference in maximum levels of pERK between KRAS^G12V^ and wild-type induced by EGFR activation, a smaller effect than observed for other active species in the pathway (Fig. 6A). Parameters that most strongly controlled the difference in maximum pERK concentrations between wild-type and mutant KRAS included rate constants for phosphorylation and dephosphorylation of ERK (*kpERK, kdpERK*) or MEK (*kpMEK, kdpMEK*) and RAF1:RAS-GTP unbinding (*kR1r*) (Fig. 6B). The reactions that involved *kdpMEK, kpMEK kdpERK,* and *kpERK* all had pERK as a reactant or a product, which made the strong effects of these parameters predictable. It was surprising, however, that the rate constants for ERK-mediated negative feedback phosphorylation of RAF1, SOS, and BRAF were excluded from the list of most sensitive parameters, as previous studies have attributed the small difference in ERK activity between wild-type and mutant to stronger ERK feedback loops in wild-type KRAS compared to mutant KRAS (15). RAF1 unbinding from RAS was a stronger determinant of pERK level difference between wild-type and mutant RAS than negative feedbacks, again highlighting RAF1 as the bottleneck for RAS mutants in the settings modeled here.

**Figure 6.**
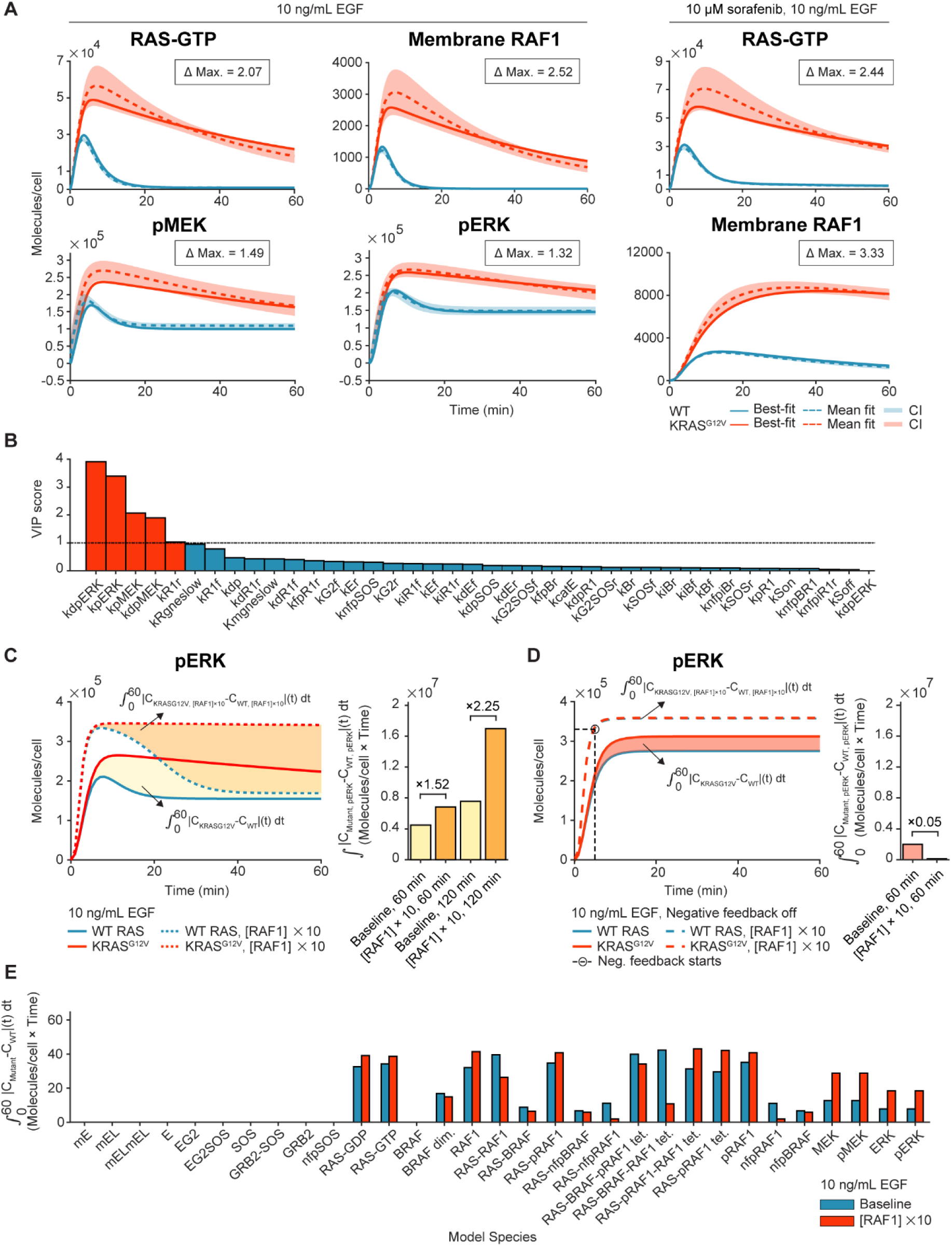
*Oncogenic KRAS^G12V^ signaling is dampened by low RAF1 abundance.* **(A)** Model- predicted dynamics of RAS-GTP, membrane-bound RAF1 with or without 10 μM sorafenib, pMEK, and pERK signaling dynamics in response to 10 ng/mL EGF for WT RAS and KRAS^G12V^ are shown. KRAS^G12V^ was simulated by a 16-fold decrease in RAS-GTP hydrolysis rate constant from its baseline value. Throughout the panels, the means of model outputs generated from the 1751 parameter sets from Fig. 3A are indicated by dotted lines, best-fit model outputs are indicated by bold lines, and 68% confidence intervals (CI) about the means are represented by shaded areas. Fold differences between maximum concentrations of indicated model species in WT and mutant dynamics (Δ max.) are boxed. **(B)** PLSR SA was performed using the 3000 uniformly sampled, random parameter sets (3000 rows × 40 columns) as independent variable matrix and predicted absolute differences in time-integrated pERK concentrations between WT RAS and KRAS^G12V^ from each parameter set (3000 rows × 1 column) were used as the dependent variable matrix. VIP scores of parameters are shown, with VIP > 1 parameters highlighted in red. **(C)** Predicted pERK dynamics of WT RAS, KRAS^G12V^ (solid, colored lines), WT RAS and KRAS^G12V^ with 10-fold increase in RAF1 expression (dashed, colored lines), simulated using the parameter values from **A**. Differences in time-integrated pERK concentrations between WT and mutant RAS with baseline RAF1 expression or with 10-fold increase in RAF1 expression are labeled and indicated by shaded areas. Fold-differences (x) between time-integrated pERK concentrations of WT and mutant RAS with baseline RAF1 expression and with 10-fold increase in RAF1 expression during 60-min or 120-min response to 10 ng/mL EGF were compared. **(D)** Predicted pERK dynamics of WT RAS and KRAS^G12V^ in the absence of negative feedback with baseline RAF1 expression (solid, colored lines) and with 10- fold increase in RAF1 expression (dashed, colored lines) generated from the best-fit from **A**. Differences in time-integrated pERK concentrations between WT and mutant RAS in the absence of feedback with baseline RAF1 expression or with 10-fold increase in RAF1 expression are labeled and indicated by shaded areas. Circle indicates the time point at which negative feedback was predicted to take effect, identified by calculating when the predicted fold difference between pERK levels in WT and mutant RAS starts becoming > 1. Fold-differences (x) between time-integrated pERK concentrations of WT and mutant RAS with baseline RAF1 expression and with 10-fold increase in RAF1 expression in the absence of feedback during 60- min response to 10 ng/mL EGF were compared. **(E)** Absolute differences in time-integrated concentrations of all model species between WT RAS and KRAS^G12V^ with baseline RAF1 expression and 10-fold increase in RAF1 expression are plotted.

To demonstrate that RAF1 acted as a bottleneck, we simulated the effect of increasing its abundance 10-fold. Compared to baseline RAF1 expression, a 10-fold increase in RAF1 expression increased the difference in time-integrated ERK activation during 60-min EGF treatment between wild-type RAS and KRAS^G12V^ by 52%, a difference that more than doubled during longer times (120 min) after EGF treatment (Fig. 6C). Perturbations in upstream pathway nodes such as RAS could be relayed to ERK only when there was sufficient (10-fold higher than baseline) RAF1 expression, and the difference became apparent only after negative feedback regulation, mainly on SOS by pERK, took effect (Fig. 6D). When the negative feedback loops were turned off, the difference in time-integrated ERK activation for wild-type versus mutant RAS decreased, and the 10-fold increase in RAF1 expression further reduced the difference by ∼100-fold, causing pERK levels of wild-type to be as high as those of mutant RAS (Fig. 6D). With the baseline RAF1 expression of 12,000 molecules/cell, the effect of constitutively active RAS on the duration of ERK signaling was most strongly dampened by RAS-pRAF1-RAF1- tetramer and RAS-pRAF1-tetramer, both of which are RAS-GTP-bound active RAF1 species (Fig. 6E). The remaining effect was distributed across RAS-GDP, RAS-GTP and other RAS-bound RAF1 and BRAF species, as they are most proximal species to the RAS-GTP hydrolysis process. On the other hand, when RAF1 expression was increased 10-fold, the effect was more heavily distributed across downstream species, such as phosphorylated MEK and ERK, suggesting that low RAF1 expression limits oncogenic RAS signaling.

### Low RAF abundance correlates with low ERK activity in human tumors and cell sensitivity to MEK inhibition

Model predictions about the effects of RAF expression were also tested by analyzing proteomics data from human LUAD and COAD tumors. In the data we analyzed, 15.6% of LUAD tumors and 16.9% of COAD tumors harbored KRAS^G12V^. RAF1 data were unavailable for LUAD, so BRAF data were used instead. In LUAD expressing wild-type or mutant KRAS, ERK1 phosphorylation on T202 and Y204 and ERK2 phosphorylation on T185 and Y187 were significantly higher in tumors with high BRAF abundance than in tumors with low BRAF abundance (Fig. 7A). In COAD, a similar effect was observed for ERK1 phosphorylation on T202 as a function of RAF1 abundance, but this relationship was not observed for ERK2 phosphorylation on T185 (Fig. 7B). No significant correlation was observed between BRAF abundance with ERK1/2 phosphorylation in wild-type or mutant KRAS in COAD. Thus, in two cancer contexts, low RAF abundance dampens ERK activity. It is worth noting that we expected to observe a larger difference made by variable RAF expression in tumors with mutant RAS than those with wild-type RAS based on our mechanistic model predictions (Fig. 6C). However, the lack of dynamic data for acute responses to growth factor in patient tumors may obfuscate the qualitative comparison to our model results.

**Figure 7.**
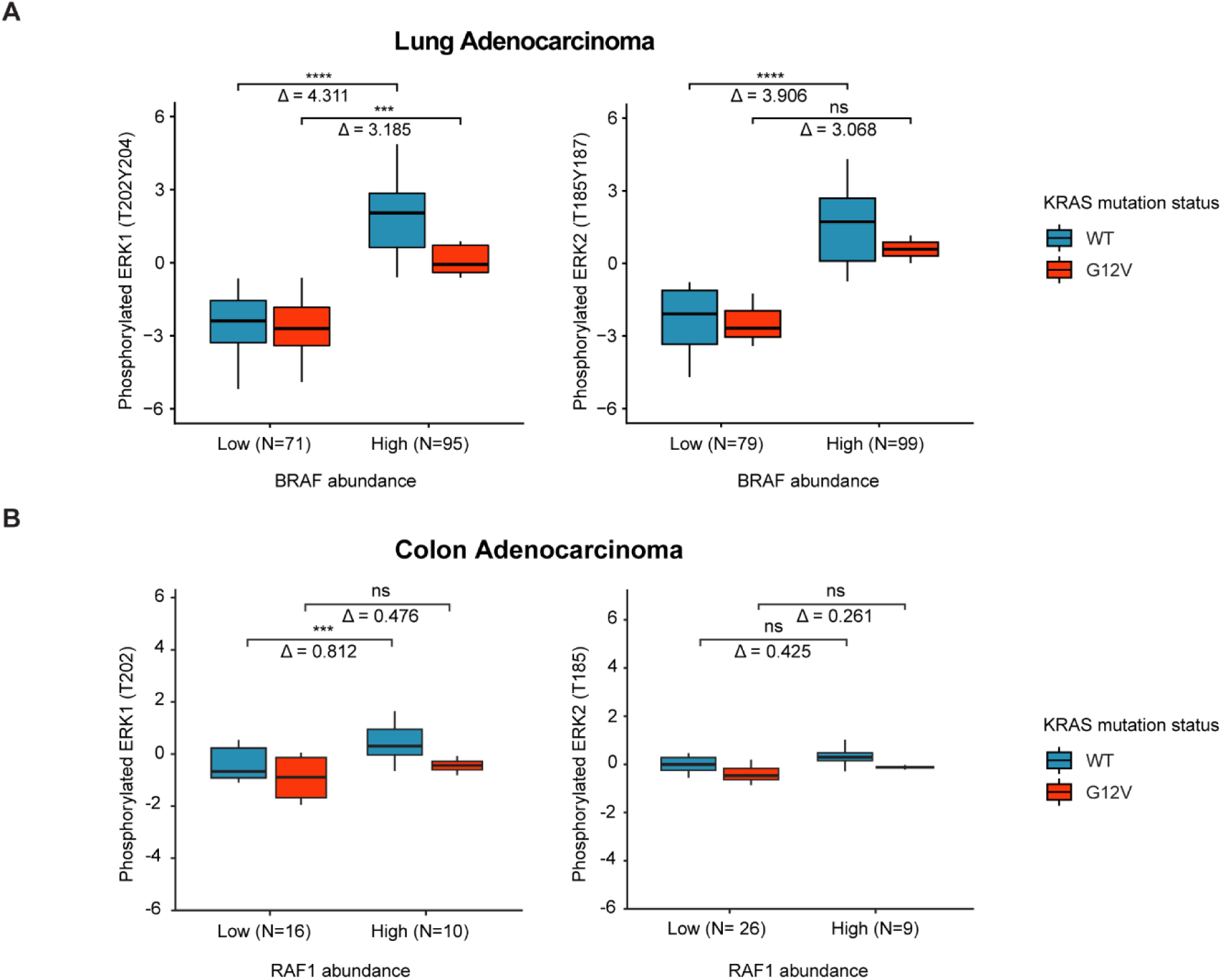
*RAF abundance is positively correlated with ERK phosphorylation in lung adenocarcinoma and colon adenocarcinoma tumors expressing wild-type KRAS and KRAS^G12V^*. **(A)** BRAF expression in CPTAC lung adenocarcinoma tumor samples expressing WT KRAS (left) and KRAS^G12V^ (right) was classified as ‘low’ or ‘high’ based on a standard median split and mapped to abundances of MAPK1 (ERK2) (top) or MAPK3 (ERK1) (bottom) phosphorylated at the indicated sites. **(B)** An analysis similar to that described in **A** was performed for RAF1 expression in CPTAC colon or adenocarcinoma tumor samples. Fold-differences in pERK1/2 for ‘low’ and ‘high’ RAF abundances are indicated, and *P* values were determined using a two-tailed Student’s *t*-test. Calculations were performed in R and Python. *, *P* < 0.05; **, *P* < 0.01; ***, *P* < 0.001; ****, *P* < 0.0001; n.s., nonsignificant.

To understand how low RAF expression affects cell phenotypes, we compared the response of CCLE cell lines with low (below 25^th^ percentile) or high (above 75^th^ percentile) RAF expression to four MEK inhibitors at their maximum tested concentrations (Fig. S4A). Raw viability data normalized by DMSO treatment (for a fixed cell seeding density) revealed that cells with low RAF expression were generally more sensitive to MEK inhibitors than cells with high RAF expression (Fig. S4B). That is, at the maximum tested inhibitor concentrations, low-RAF cells exhibited greater overall reductions in cell numbers than did high-RAF cells. At the same time, the half-maximal inhibitory concentrations (IC50) of the MEK inhibitors for the proliferation phenotype were unaffected by RAF expression (Fig. S4C), indicating RAF abundance did not impact the fraction of total response possible to increases in drug abundance. Both observations can be true simultaneously because the absolute change in viability effected by a drug is distinct from the rate at which the total possible drug effect manifests as drug concentration increases. Curiously, when we simulated MEK inhibition in the setting of response to EGF by simply reducing the initial MEK abundance by up to 100% (a choice that obviated the need to otherwise introduce additional model species and rate constants for direct simulation of MEK inhibition), we observed that the predicted time-integrated ERK activity was more sensitive to increases in MEK inhibition for low RAF abundance (Fig. S4D). Thus, while the model predicts that ERK is more responsive to MEK inhibition in a low-RAF context, that biochemical effect (if it indeed occurs in cells) may be insufficient to drive a viability phenotype dose-response effect.

### RAF1 stochasticity results from low RAF1 abundance and processes upstream of RAF

The low abundance of RAF1 at the membrane measured in prior work (9) (∼200 RAF1 molecules per cell at 6-8 min after EGFR activation) raises a question of whether certain levels of the EGFR-ERK pathway operate in a stochastic fashion in RAF-bottleneck settings and how system responses to perturbations may be affected by stochastic homotypic RAF1 interactions or heterotypic RAF1/RAS or RAF1/BRAF interactions. There is precedent for this notion. For example, qualitative differences in cell motility arise due to stochastic signaling dynamics for Rho GTPases expressed at 2000-3000 molecules per cell (66). To begin to address these issues, we simulated the response of GTP-RAS, membranous RAF1, pMEK, and pERK to EGF using a stochastic solver for five different random seeds. Random seeds are used to generate reaction rates between species sampled from an exponential distribution (48), with the seed- generating function set so that the mean of the simulated distribution converges to the theoretical rates predicted by Michaelis-Menten or first-order mass action (67). Julia was used for these calculations because the platform allows parallelization of stochastic simulations across as many parameter sets as desired. As expected, dynamics were more stochastic at the level of membranous RAF1 than at other pathway nodes, but the effect was modest with deviations among seeds that were small compared to the magnitude of the signal (Fig. 8A).

**Figure 8.**
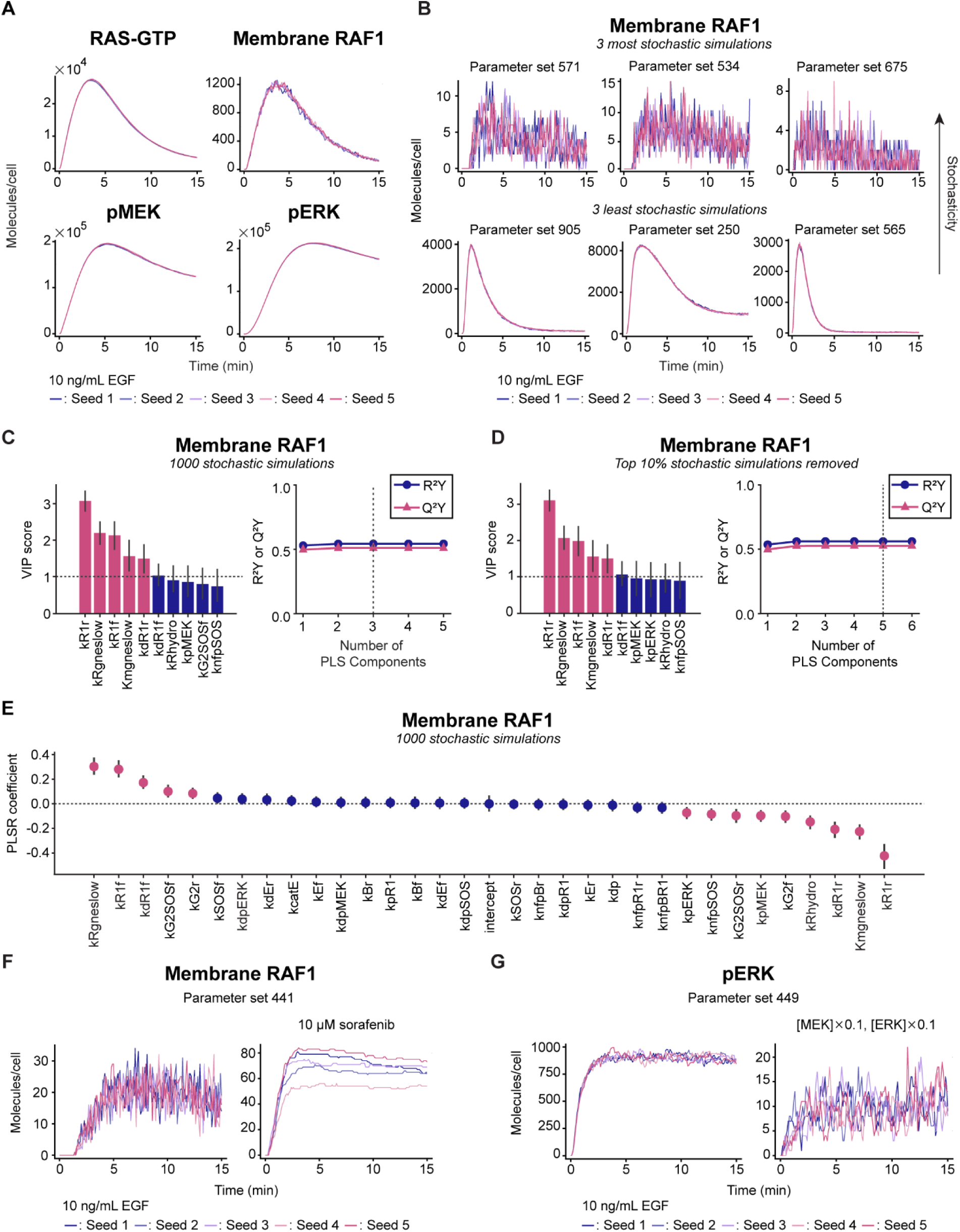
*Stochasticity arising from low membrane RAF1 abundance is most strongly controlled by RAF1:RAS-GTP interactions*. **(A)** Stochastic model predictions of RAS-GTP, membrane RAF1, pMEK, and pERK concentrations in response to 10 ng/mL EGF with the best-fit parameter values from Fig. 3A (N=5 differentially seeded, stochastic runs). **(B)** Stochastic membrane RAF1 dynamics with the three parameter sets that produced the highest (top) or lowest (bottom) membrane RAF1 stochasticity scores, as defined in *Results*, out of 1000 parameter sets. **(C)** PLSR SA was performed using 1000 LHS parameter sets (1000 rows × 40 columns) as the independent variable matrix, and membrane RAF1 stochasticity scores for each parameter set (1000 rows × 1 column) were used as dependent variable matrix. The top 10 parameters ranked by PLSR SA VIP scores are shown. Confidence intervals for the VIP scores were obtained by running the PLSR SA model 10,000 times by random sampling of the X and Y matrices with resampling. Error bars indicate 95% confidence intervals. Parameters highlighted in pink have lower bounds of the confidence intervals of VIP scores > 1. **(D)** PLSR SA with the top 10% of stochastic simulations with the highest stochasticity scores removed. Error bars indicate 95% confidence intervals. Parameters highlighted in pink have lower bounds of the confidence intervals of VIP > 1. **(E)** PLSR coefficients from bootstrapped PLSR SA described in **C** are shown. Dots indicate means of 10,000 bootstrapped PLSR coefficient samples, and bars indicate 95% confidence intervals. Parameters with mean PLSR coefficients highlighted in pink have nonzero lower and upper bounds of the confidence intervals of the coefficients. **(F)** Stochastic membrane RAF1 dynamics in the absence (left) or presence of 10 μM sorafenib (right) with the parameter set that produced the most stochastic behavior both with and without sorafenib. **(G)** Stochastic pERK dynamics with the parameter set that produced the highest membrane RAF1 stochasticity score with baseline protein expression levels (left) or with 10-fold decreases in MEK and ERK expression (right). All calculations were performed in Julia.

To identify conditions that would promote even more RAF1 stochasticity at the membrane, we generated 1000 parameter sets using LHS wherein individual parameters were perturbed by factors ≤ 10 above or below baseline values. To quantify the effects across different parameter sets, we defined a *stochasticity score* as the variance in time-integrated membrane RAF1 dynamics among the five differentially seeded simulations. Certain parameter sets resulted in substantial membrane RAF1 stochasticity, while others did not, as was apparent in the three most- or least-stochastic simulations run (Fig. 8B). PLSR SA with the 1000 LHS parameter sets from Fig. 8B as independent variables and the corresponding stochasticity scores as the dependent variables revealed that the parameters that most strongly controlled stochastic membrane RAF1 levels were similar, but not identical, to those that controlled deterministic membrane RAF1 solutions shown in Fig. S3A,B (Fig. 8C). Similar results were obtained when ∼200 stochastic simulations were compared to deterministic simulations in VCell (Fig. S5A-C). Notably, parameters controlling phosphorylation of ERK and MEK (*k_pERK_, k_pMEK_*) decreased in importance in the control of stochastic membrane RAF1 solutions relative to what was observed for the deterministic simulations, suggesting that MEK and ERK do not control membrane RAF1 stochasticity. To confirm this, we rebuilt the PLSR SA model after removing the top 10% of the most stochastic solutions. RAS- and RAF1-proximal parameters were identified to control maximum membrane RAF1 levels predicted by the remaining stochastic solutions, suggesting that stochasticity played a negligible role in maximal RAF1 activation (Fig. 8D).

Plotting stochasticity scores over ranges of strongly controlling parameters provided additional insight. To obtain contrasting cases of two-dimensional parameter spaces with high versus low influence on membrane RAF1 stochasticity, we selected a subset of parameters based on PLSR SA for stochasticity scores as the dependent variable (Fig. 8E). PLSR SA of membrane RAF1 stochasticity scores against 1000 LHS-sampled parameter sets revealed that the same rate constants that controlled the EGFR-ERK system identified by PLSR SA with filtered parameter sets and Sloppiness analysis (Fig. S1B), namely the RAF1:RAS-GTP (un)binding at the membrane and RAS activation rates, were also the most important predictors of stochasticity. When interpolated stochasticity scores were plotted as a function of the parameter values, parameter values that make RAF1 binding from RAS unlikely significantly increased membrane RAF1 stochasticity while the parameters that controlled stochasticity scores the least produced a flat contour (Fig. S5D). The stochastic model also predicted that membrane RAF1 in the presence of sorafenib could have qualitatively different behaviors depending on the differential rates of RAF1 binding to RAS-GTP (i.e., the different seeds) (Fig. 8F). This difference is manifested in form of divergence in trajectories among random seeds, which is distinct from noisy, local oscillations in stochastic simulation results in the absence of sorafenib.

To determine if the effects of membrane RAF1 stochasticity could propagate within the pathway, we selected a parameter set with one of the top stochasticity scores and inspected the degree of predicted stochasticity at the level of pERK. Predicted stochasticity was modest for baseline protein expression levels, but increased substantially when the expression of MEK and ERK were decreased by just ten-fold (Fig. 8G), leaving them in the tens-of-thousands of copies per cell and well within the range of expression values observed in some cell lines, including HMEC breast epithelial cells and HS578T breast cancer cells (4). Decreased MEK and ERK expression did not influence membrane RAF1 levels (Fig. S5E). Thus, these model calculations suggest that stochasticity at the level of membrane RAF1 could indeed plausibly propagate to ERK in certain settings.

## DISCUSSION

Our results suggest that RAF is frequently a strong regulator of the ERK pathway downstream of EGFR by acting as a low-abundance stoichiometric bottleneck and limiting overall pathway flux. Low RAF abundance effects on signaling impact cancer cell phenotypes, demonstrated by significant absolute differences in the viability response of cancer cells with low or high RAF expression to MEK inhibitors. RAF is not typically recognized as playing this role, but prior work (4,9) and the analysis shown here of common cell lines and patient tumor data suggest that the low-RAF abundance setting is reasonably common. In principle, any pathway intermediate can be stoichiometrically limiting. In normal breast cells and breast cancer cells, for example, SOS and GAB1 can be even less well expressed than RAF1 and may be rate-limiting for ERK activation (4). Consistent with this notion, our model predicted that when RAF1 stoichiometric limitations are relieved, GRB2/SOS interactions and SOS abundance regulate peak ERK activation more strongly than do RAF-related parameters. For stoichiometric settings such as those encountered in HeLa cells, sensitivity and Sloppiness analysis revealed that EGFR-ERK signaling (at six nodes spread throughout the pathway) was most sensitively regulated at the level of RAF1 even with an overall SOS abundance (7700 molecules/cell) that was lower than that for RAF1 (12,000 molecules/cell). This occurred in our simulations in part because translocation to the membrane was much less efficient for RAF1 (10-15%) than for SOS (55%), and binding affinity of RAF1 at the membrane was ∼79 fold lower than that of SOS. The extremely low tendency for RAF1 to membrane-localize makes it unique as a low-abundance pathway intermediate.

The model is especially complex in the vicinity of RAS and RAF due to numerous binding and reaction terms. One could wonder if the sheer number of parameters in this region of the model is somehow responsible for the predicted importance of RAF1 in governing system behavior. We can attribute the predicted importance of RAS- and RAF1-related parameters to the low RAF1 expression with confidence, however, because increased RAF1 expression reduced the predicted importance of RAS- and RAF1-related parameters in favor of parameters that more proximally controlled ERK itself. Moreover, when only ERK activity was considered as the model output, parameters for MEK and ERK activation became more important for determining system output, with the parameter describing RAF1 unbinding from RAS-GTP as secondarily important. This is qualitatively consistent with a prior study showing that control over transient ERK activity was distributed among RAF and MEK/ERK phosphorylation/dephosphorylation rates in HeLa cells (68).

Stochastic behavior in signal transduction is typically considered, if at all, at the transcriptional level. Many transcription factors are expressed at low copy numbers (e.g., as low as 5 copies/cell for c-Fos (69)), and their intrinsic fluctuations act as a source of noise to result in phenotypic variability (70). However, our results suggest that stochasticity in the EGFR-ERK pathway can arise upstream of transcription factors, consistent with the conclusion reached by an ODE-based model of the tumor necrosis factor pathway that attributed heterogenous cell viability to varying protein levels of BID, BAX, and BCL2 (71). The model prediction that low RAF1 abundance leads to intrinsic stochasticity (non-deterministic system behavior) at specific nodes of the EGFR-ERK signaling cascade may explain why some experimental measurements are persistently technically noisy or irreproducible. For example, our results suggest that the experimentally observed difference in membrane RAF1 variance with or without sorafenib (9) (lower variance with sorafenib) could have arisen from the greater abundance of RAF1 at the membrane for cells treated with sorafenib versus untreated (9). Our results also show how stochasticity caused by low RAF1 abundance can be smoothed at downstream nodes when those species are more abundant or increasingly propagated when downstream species are also expressed with low abundance. Stochasticity effects also impacted the results of overall sensitivity analysis.

In discussing stochasticity effects in signaling, care must be used to specify the source and type. For example, a previously developed classification model predicted that fluctuations in RAF1 and BRAF expressions engendered cell-to-cell variation in ERK activity and proliferation among MCF10A cells (72). While this result may appear to stand in contrast to our finding that stochasticity effects for low-abundance RAF1 do not propagate to ERK, the issue lies in the type of stochasticity being described. Because RAF1 and BRAF are critical determinants of ERK signaling (consistent with our inferences), cell-to-cell variation in their expression (termed stochasticity by the authors of that paper) leads to cell-to-cell variability in ERK activity. That can be true even without the effects of intrinsic stochasticity or non-deterministic system behavior, wherein cells with identical expression levels of the same pathway intermediates have different reaction trajectories simply because the protein abundance is too low for deterministic behaviors.

Comparison of the influential system parameters nominated by our analysis with those identified by others should be made with some caution, in part because our sensitivity analysis considered six pathway nodes simultaneously for a holistic assessment of system performance. However, the primary output of interest in this system is typically phosphorylated ERK (19,68,73,74). We considered model outputs at multiple pathway nodes because many more parameter combinations are likely to be able to explain pERK dynamics alone than are capable of explaining system dynamics at all six nodes we considered simultaneously. Thus, our approach is more likely to lead to a model that is predictive of network-level scenarios not considered in model training. While some well-known regulatory modes, including induced expression of dual specificity phosphatases, receptor activation/deactivation cycling, and receptor dimerization (7,19,74,75), have previously been identified as strong regulators of system-wide control, our sensitivity analysis points to RAF1:RAS-GTP binding as the stronger determinant of overall system behavior in the context of low RAF1 abundance. In our model, the outputs of peak and time-integrated ERK activity are secondarily sensitive to changes in other receptor- and adaptor-level rate constants and expression levels (e.g., GRB-SOS binding, receptor dimerization, and SOS concentration), in qualitative agreement with other reports (4,7,74–76).

Our understanding of how RAF1 abundance affects EGFR-ERK pathway dynamics may also be impacted by model assumptions and technical limitations. For example, replacing our typical assumption of first-order enzyme-catalyzed kinetics by saturable Michaelis-Menten kinetics may more realistically constrain reaction rates, as in Futran *et al.* (77), but would simultaneously increase the number of species and rate constants. As another example, in the stochasticity sensitivity analysis we performed, the degree of stochasticity would be more generalizable if it included extrinsic factors such as diffusion through use of the Gillespie algorithm with more simulations and incorporation of intrinsic (e.g., intermolecular collisions) and extrinsic (e.g., cell cycle effects) fluctuations (78). However, the increased complexity of a reaction-diffusion model with the advanced stochastic solver comes at the cost of substantially increased computation time (79). In the future, as it becomes more computationally feasible to address these issues in a system of the size considered here, it will be important to assess the impact of those types of assumptions and limitations on qualitative conclusions.

To complete our analysis, we used several different computing platforms due to issues of speed, solver availability, and parallel computing capabilities, but many of the obstacles that led us to integrate multiple platforms can now be overcome by the diverse packages available in the Julia programming language. Some relevant Julia packages were still in development when we began the study. While we did not use Julia throughout this work, it served as a useful tool to extend the model analyses performed in VCell and MATLAB. Particular benefits of Julia are the simplicity of encoding a reaction network, speed of the ODE solver, and the freedom to choose how to distribute memory among large sets of parameter values. These capabilities promise opportunities for beginner systems biologists to conduct advanced sensitivity analyses without the need to learn multiple programming languages or software (80).

Finally, our comparison of traditional sensitivity analyses and model Sloppiness analysis yielded previously unappreciated similarities and differences between parameters that control uncertainty versus sensitivity of model outputs of EGFR-ERK signaling with low RAF1 abundance. Sloppiness analysis and sensitivity analysis with respect to dynamics of the key pathway nodes used in model training identified RAS activation and RAF1:RAS-GTP (un)binding rate constants as parameters that should be measured accurately to build a good model that captures the observed signaling behaviors. On the other hand, sensitivity analysis with respect to maximal ERK activity identified MEK and ERK (de)phosphorylation parameters as important determinants of ERK activity in the presence of large perturbations. This highlights the distinct utility of sensitivity analysis serves as a hypothesis generator versus model Sloppiness analysis as a tool for model calibration.

## AUTHOR CONTRIBUTIONS

**S.H. Lee**: Conceptualization, data curation, formal analysis, investigation, visualization, methodology, writing–original draft, writing–review and editing. **P.J. Myers**: Conceptualization, formal analysis, visualization, methodology, writing–original draft, writing–review and editing. **K.S. Brown**: Investigation, methodology, funding acquisition, writing–review and editing. **L.M. Loew**: Investigation, funding acquisition, writing–review and editing. **A. Sorkin**: Investigation, data curation, funding acquisition, writing–review and editing. **M. J. Lazzara**: Conceptualization, supervision, funding acquisition, project administration, writing–original draft, writing–review and editing.

## ACKNOWLEDGEMENTS

This work was supported by NSF MCB 1716537 (MJL), 1715132 (AS), 1716075 (LL), 1715342 (KB), NIH U54 CA274499 (MJL), NIH R35 GM148363 (AS) and the University of Virginia Biomedical Data Sciences training grant NIH T32LM012416. The Virtual Cell is supported by NIH R24 GM137787. The authors acknowledge Research Computing at the University of Virginia for computational resources and technical support. Figure 2 was created with BioRender.com.

## SUPPORTING MATERIAL

**Figure S1.**
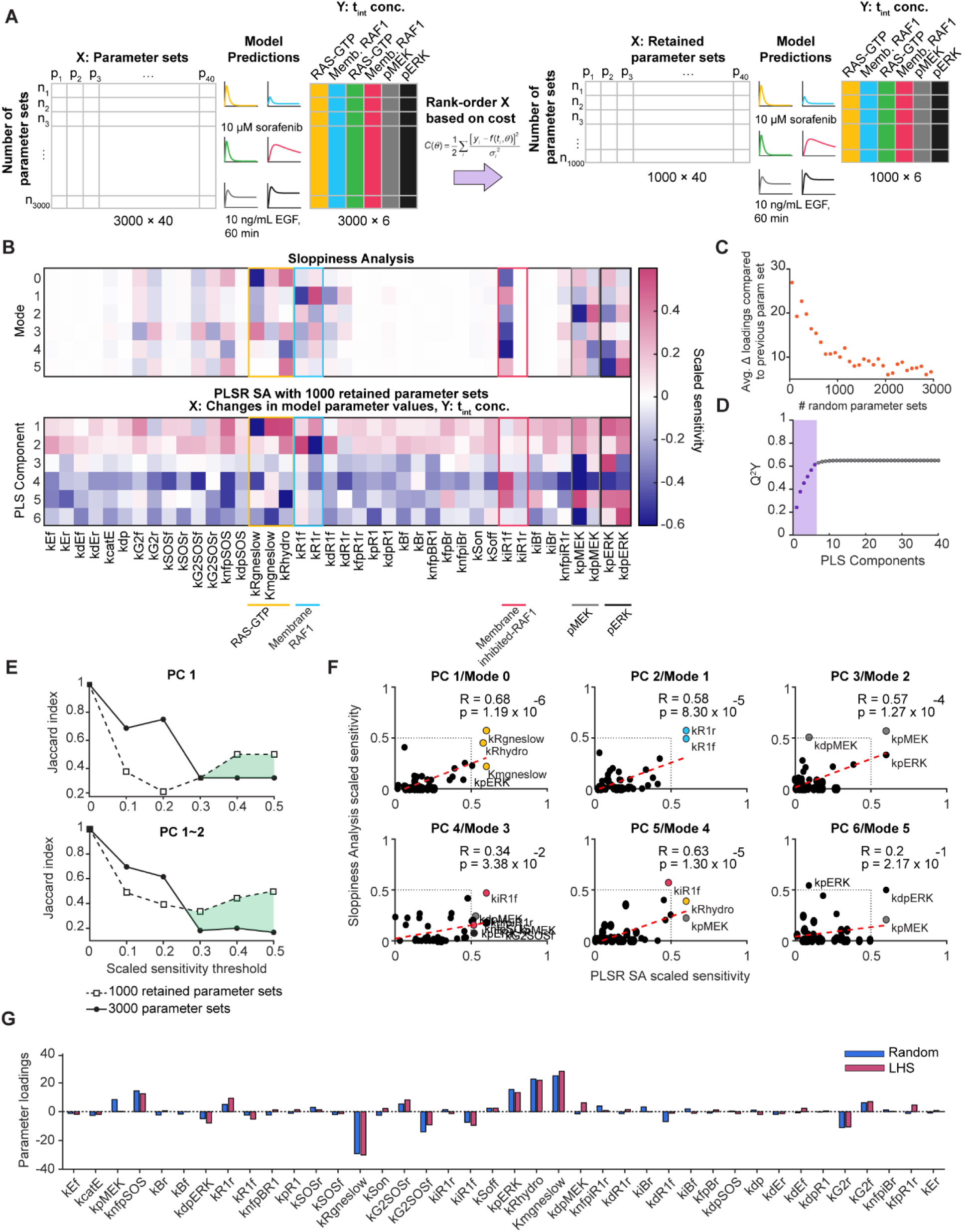
There is greater overlap between multivariate sensitivity and Sloppiness analyses when parameter sets that produce relatively large deviation from experimental data are removed. **(A)** Workflow of filtering parameter sets based on the cost associated with model predictions obtained by each parameter set. The 3000×40 matrix used to build PLSR SA in Fig. 4 was reduced to a 1000×40 matrix after retaining just one-third of the parameter sets that produced the lowest cost. The retained parameter sets and corresponding model-predicted, time-integrated concentrations of RAS-GTP and membrane RAF1 with or without sorafenib, pMEK, and pERK in response to 10 ng/mL EGF during 60 min were used for a new PLSR SA to compare with Sloppiness analysis. **(B)** Projection of parameters onto the first six modes from model Sloppiness analysis and parameter loadings in the first six PLS components from PLSR SA of model-predicted, time-integrated concentrations of GTP-RAS, membrane RAF1, pMEK, and pERK against the 1000 retained parameter sets were compared. PLSR SA parameter loadings were scaled to the range of parameter projections on the FIM in Sloppiness analysis (- 0.6∼0.6) and plotted as scaled sensitivities. **(C)** Average differences (Δ) in all parameter loadings between PLSR SA models built with different numbers of random parameter sets using the maximum membrane RAF1 concentrations as the dependent variable. **(D)** Predictive power determined by cross-validation (Q^2^Y) of PLSR SA model with filtered parameter sets for increasing numbers of PLS components are plotted. **(E)** Jaccard indices of shared parameters that had scaled sensitivities above the indicated threshold values in the first component/mode (top) and the first two components/modes combined (bottom) in PLSR SA and Sloppiness analysis are shown. The indices were calculated for PLSR SA with 3000 and 1000 retained parameter sets. **(F)** Scatter plots of scaled sensitivities in Sloppiness analysis and PLSR SA are shown. Pearson’s correlation coefficients (R) of scaled sensitivities are calculated for each respective component and mode. Parameters with scaled sensitivity from both analyses > 0.5 are labeled and filled with color. *P* values were calculated using one-tailed Student’s *t*-test. **(G)** PLSR SA parameter loadings using random and LHS-sampled parameter sets as independent variable matrix and maximum membrane RAF1 levels as dependent variable matrix are compared. All calculations were performed in MATLAB.

**Figure S2.**
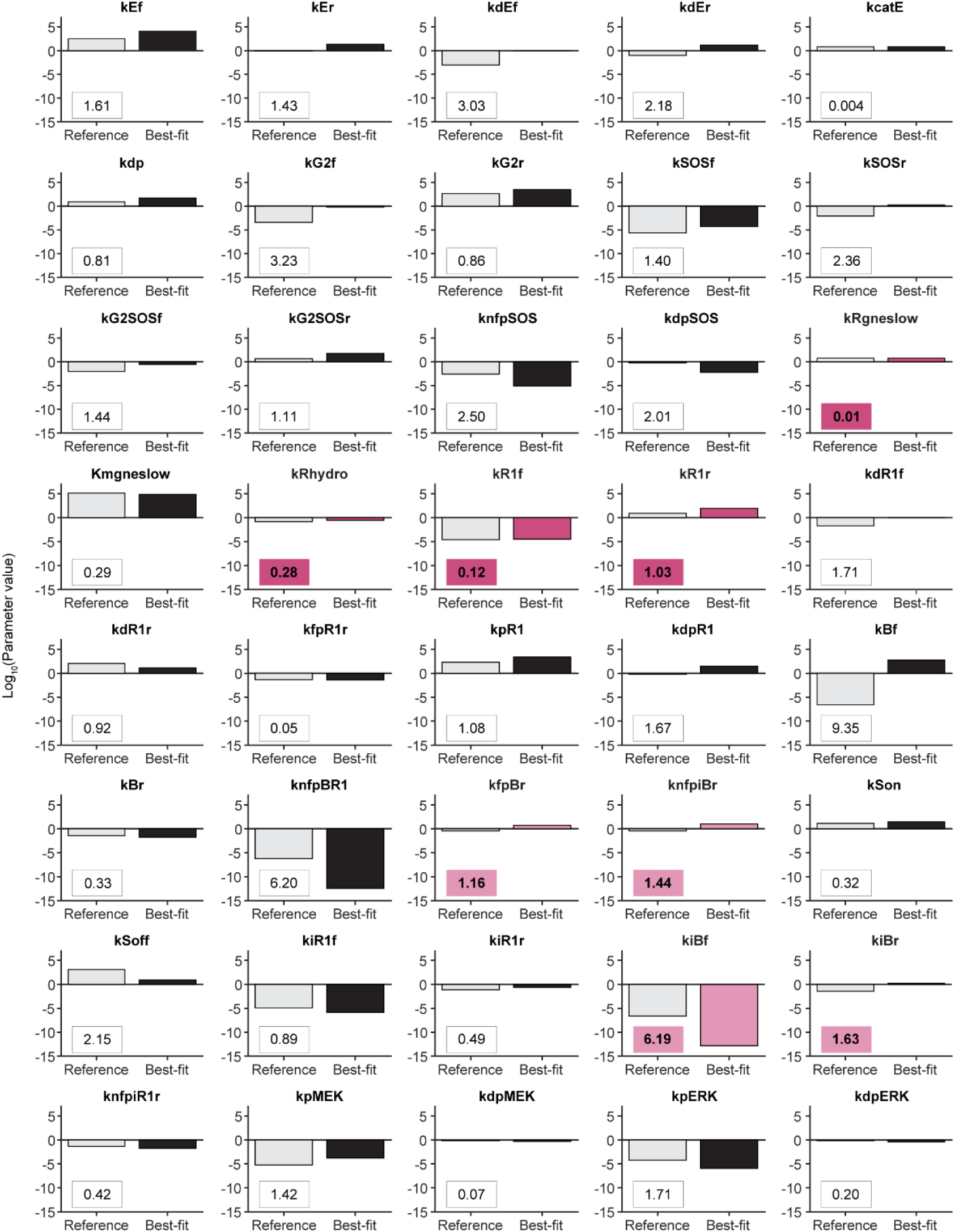
*Fitted and reference values closely match for stiff parameters identified by Sloppiness analysis.* Best-fit parameter values and reference values from Tables S4 and S7, respectively, are plotted, with their log-fold differences boxed. The four stiffest parameters from modes 0 and 1 (Sloppiness analysis in Fig. 4) are highlighted in dark pink, and the four sloppiest parameters from modes 0 and 1 are highlighted in light pink.

**Figure S3.**
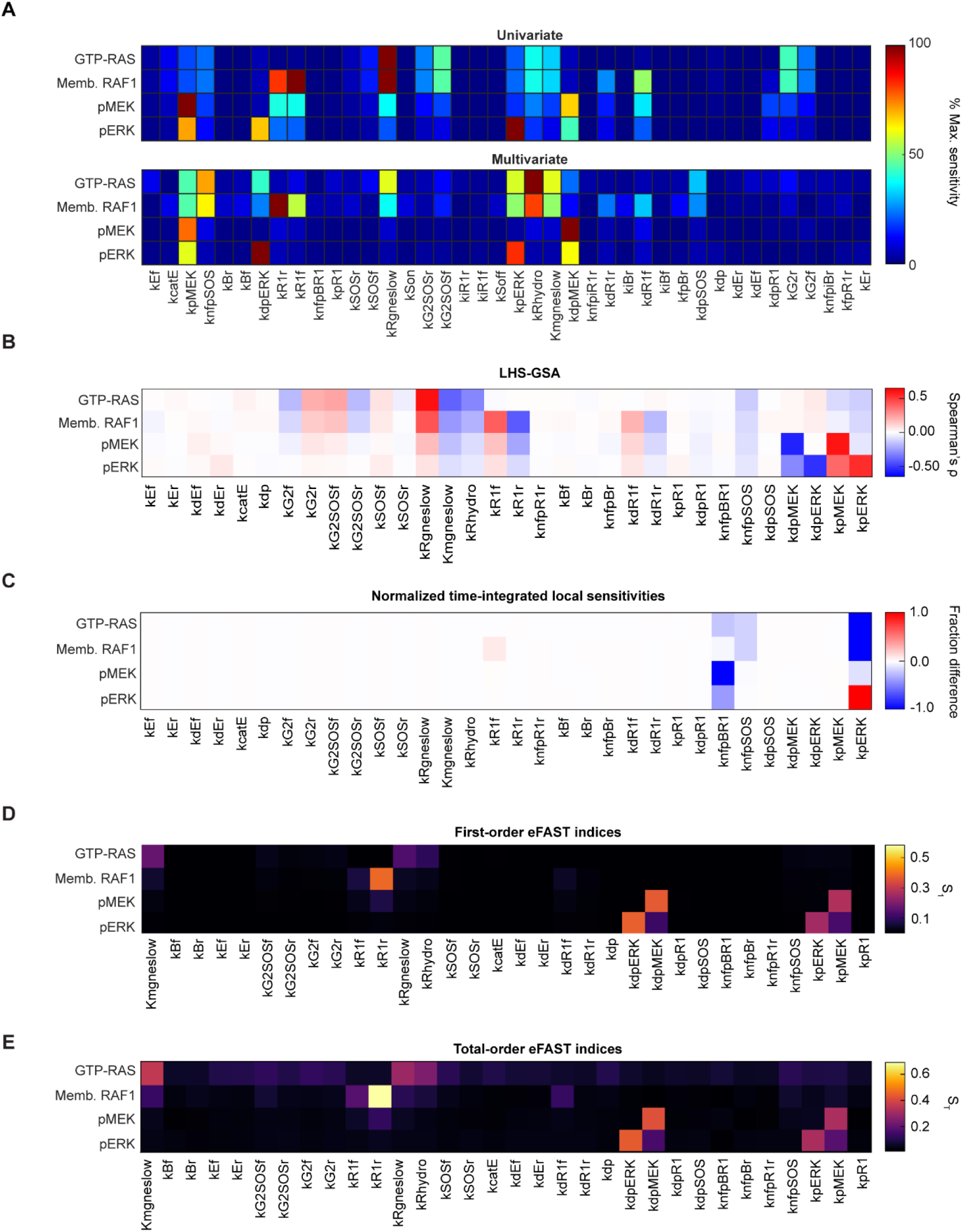
*Local and univariate sensitivity analyses identify fewer influential parameters than multivariate approaches.* **(A)** Univariate sensitivity analysis was conducted by taking the percent difference in the predicted maximum RAS-GTP, membrane-bound RAF1, pMEK, and pERK concentrations in response to 10 ng/mL EGF as a result of increasing and decreasing individual parameter values by a factor of 10. Multivariate sensitivity analysis was conducted by taking the regression coefficients from PLSR SA against 3000 randomly sampled parameter sets based on log-uniform distribution. Only the model outputs to which the model was fitted were chosen for sensitivity analyses. **(B)** Spearman’s correlation coefficients (ρ) of changes in the maximum total RAS-GTP, membrane RAF1, pMEK and pERK concentrations and LHS-sampled parameter values. **(C)** Local sensitivities of time-integrated total RAS-GTP, membrane RAF1, pMEK, and pERK concentrations in the absence of sorafenib, normalized to the maximum local sensitivity value for each model output. **(D)** First-order eFAST index (S1) of each parameter calculated for the maximum total RAS-GTP, membrane RAF1, pMEK and pERK concentrations, normalized to the maximum first-order eFAST index for each model output. **(E)** Total-order eFAST index (ST) calculated for the maximum total RAS-GTP, membrane RAF1, pMEK and pERK concentrations as model outputs. Univariate and multivariate SA were performed in MATLAB, local SA, LHS- GSA and eFAST analyses were performed in Julia.

**Figure S4.**
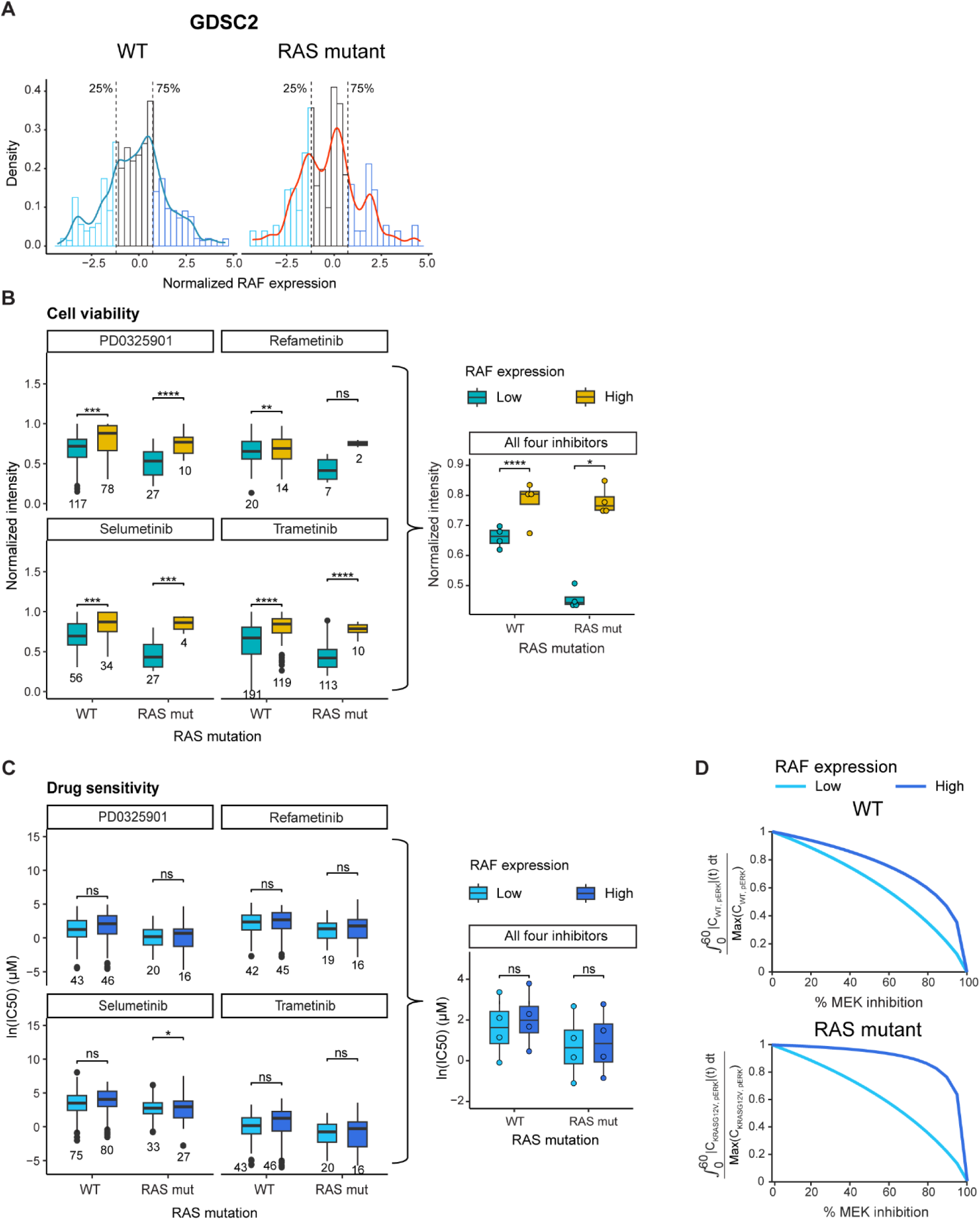
*Low RAF expression sensitizes cancer cells to MEK inhibitors*. **(A)** GDSC2 cell viability data were combined with CCLE proteomics data to create a data frame including RAF protein abundance (normalized as reported for CCLE mass spectrometry) and RAS mutational status for 239 cell lines. Data are displayed as a histogram (displayed as density) with the kernel density function overlayed for the normalized RAF expression. Dotted lines indicate the cutoffs for cellular RAF protein expression in the lowest 25% (Low) or highest 25% (High). **(B)** Normalized cell viability fluorescence intensity measurements (CellTiterGlo assay) indicating response to MEK inhibitors at the maximum concentrations tested for each: PD0325901 (2.5 μM), refametinib (10 μM), selumetinib (10 μM), and trametinib (1 μM). The reported fluorescence intensities were normalized to fall within a range bounded by measurements for DMSO (control) and blank wells and were compared for low or high RAF expression. The plot at right shows the same results aggregated over all four inhibitors. **(C)** Natural log-transformed IC50 values reported in viability data described in **B** for PD0325901, refametinib, selumetinib, and trametinib were compared between cell lines with low or high RAF expression. The plot at right shows the same results aggregated over all four inhibitors. **(D)** The EGFR-ERK model developed here was used to predict the effects of increasing RAF1 expression by 10-fold. To simulate MEK inhibition, the initial concentration of MEK was reduced by up to 100% of its baseline value. Simulations were run for wild-type RAS and the KRAS^G12V^ mutant, which was modeled through a 16-fold decrease in RAS-GTP hydrolysis rate constant from the baseline value.

**Figure S5.**
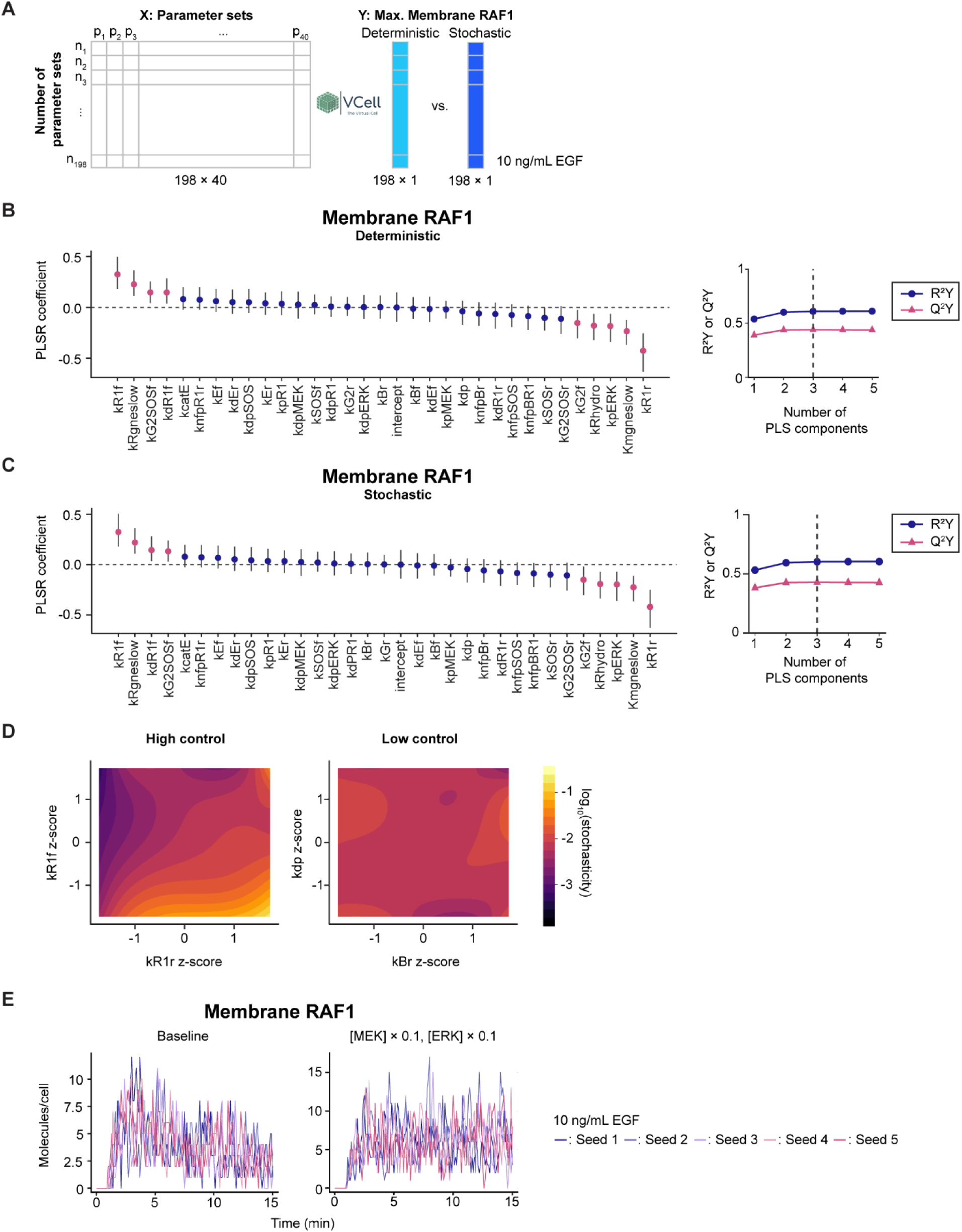
*Deterministic and stochastic membrane RAF1 levels are most strongly controlled by RAS:RAF1 binding parameters.* **(A)** Summary of PLSR SA based on VCell simulations. Independent variable matrix (X) was set to 198 (n) sets of mean-centered, variance-scaled, LHS-sampled model parameter (p) values, and the dependent variable matrix (Y) was set to deterministic or stochastic model-predicted maximum membrane RAF1 concentrations in response to 10 ng/mL EGF. **(B)** PLSR coefficients for each parameter using the deterministic solver in VCell (left), cumulative percent variance explained (R^2^Y), and percent variance predicted (Q^2^Y) of each component in the PLSR model are shown (right). Confidence intervals for the PLSR coefficients were obtained by running the PLSR SA model 10,000 times by random sampling of the X and Y matrices with resampling. Error bars indicate 95% confidence intervals. Parameters with mean PLSR coefficients highlighted in pink have nonzero lower and upper bounds of the confidence intervals of the coefficients. **(C)** PLSR coefficients with LHS- sampled parameter sets regressed against 198 stochastic batch simulations from VCell, cumulative percent variance explained (R^2^Y) and percent variance predicted (Q^2^Y) of each component in the PLSR model are shown (right). Parameters with mean PLSR coefficients highlighted in pink have nonzero lower and upper bounds of the confidence intervals of the coefficients, obtained by the method described in **B**. **(D)** Contour plots of *z*-scored parameter values of the parameter pairs with the highest absolute regression coefficients nominated from Fig. 8E mapped against interpolated stochasticity. Color bar indicates log-scaled stochasticity scores. **(E)** Stochastic membrane RAF1 dynamics predicted with the parameter set that produces the most stochastic membrane RAF1 dynamics with baseline protein expressions (Left) and with 10-fold decrease in MEK and ERK expressions (Right).

**Table S1.** Software and algorithms.

| Resource | Identifier | Source or reference |
| --- | --- | --- |
| Virtual Cell v7.4.0 | https://vcell.org/ | (37,38) |
| R v4.2.2 | https://www.r-project.org/ | *R Core Team* |
| factoextra v1.0.7 (R package) | https://doi.org/10.32614/CRAN.package.factoextra | Alboukadel Kassambara, Fabian Mundt |
| FactoMineR v2.11 (R package) | https://doi.org/10.32614/CRAN.package.FactoMineR | (81) |
| cluster v2.1.4 (R package) | https://doi.org/10.32614/CRAN.package.cluster | (82) |
| ggplot2 v3.4.2 (R package) | https://doi.org/10.32614/CRAN.package.ggplot2 | (81,83) |
| reticulate v1.30 (R package) | https://rstudio.github.io/reticulate/ | Kevin Ushey, JJ Allaire, Yuan Tang |
| gdscIC50 (R package) | https://github.com/CancerRxGene/gdscIC50/tree/master | (84) |
| SloppyCell 2017 (Python v2 package) | https://sloppycell.sourceforge.net/ | (39,83,85) |
| cptac v1.1.2 (Python v3 package) | https://pypi.org/project/cptac/ | (86) |
| MATLAB R2020a | https://www.mathworks.com | The MathWorks, Natick, MA |
| fminsearchbnd (MATLAB function) | https://www.mathworks.com/matlabcentral/fileexchange/8277-fminsearchbnd-fminsearchcon | John D'Errico (2024). fminsearchbnd, fminsearchcon, MATLAB Central File Exchange. |
| LHS_Call (MATLAB function) | http://malthus.micro.med.umich.edu/lab/usanalysis.html | (26) |
| labelpoints v4.1 (MATLAB function) | http://www.mathworks.com/matlabcentral/fileexchange/46891-labelpoints | Adam Danz (2024). labelpoints, MATLAB Central File Exchange. |
| Julia v1.10.0 | https://julialang.org/ | (26,41) |
| Catalyst.jl v13.5.1 | https://docs.sciml.ai/Catalyst/stable/ | (42) |
| ModelingToolkit.jl v8.73.2 | https://docs.sciml.ai/ModelingToolkit/dev/ | (41,43) |
| DifferentialEquations.jl v7.11.0 | https://docs.sciml.ai/DiffEqDocs/stable/ | (42,44) |
| MLJ.jl v0.20.2 | https://juliaai.github.io/MLJ.jl/stable/ | (43,87) |
| SciMLSensitivity.jl v7.51.0 | https://docs.sciml.ai/SciMLSensitivity/stable/ | (44,88) |
| GlobalSensitivity.jl v2.4.0 | https://docs.sciml.ai/GlobalSensitivity/stable/ | (50,87) |

**Table S2.**
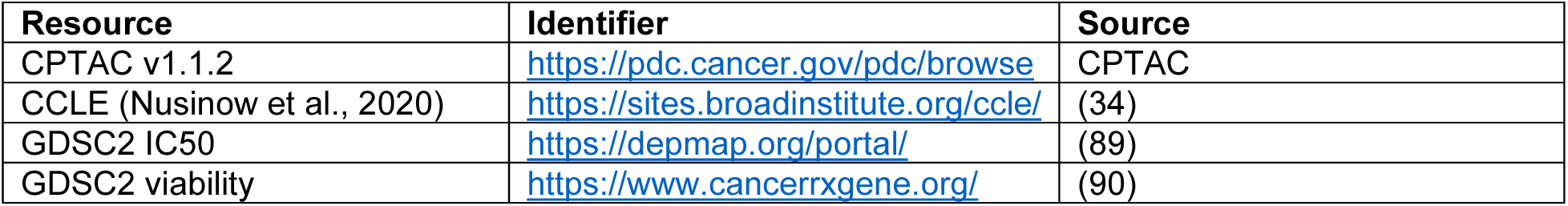
Datasets.

| Resource | Identifier | Source |
| --- | --- | --- |
| CPTAC v1.1.2 | https://pdc.cancer.gov/pdc/browse | CPTAC |
| CCLE (Nusinow et al., 2020) | https://sites.broadinstitute.org/ccle/ | (34) |
| GDSC2 IC50 | https://depmap.org/portal/ | (89) |
| GDSC2 viability | https://www.cancerrxgene.org/ | (90) |

**Table S3.** Species labels and names in the EGFR-ERK model.

| **Label** | **Species** |
| --- | --- |
| mE | EGFR monomer |
| mEL | EGF-bound EGFR monomer |
| mELmEL | EGF-bound EGFR dimer |
| E | EGF-bound pEGFR dimer |
| SOS | SOS1/2 |
| GRB2_SOS | GRB2 bound to SOS |
| EG2 | EGF-bound pEGFR dimer bound to GRB2 |
| EG2SOS | EGF-bound pEGFR dimer bound to SOS |
| RAS_GDP | GDP-bound RAS |
| RAS_GTP | GTP-bound RAS |
| BRAF | BRAF monomer |
| BRAF_dimer | BRAF dimer |
| iBRAF | BRAF inhibited by sorafenib |
| iBRAF_dimer | Inhibited BRAF dimer |
| BRAF_iBRAF_dimer | BRAF-inhibited BRAF dimer |
| RAF1 | RAF1 |
| pRAF1 | Phosphorylated RAF1 |
| iRAF1 | RAF1 inhibited by sorafenib |
| Ras_RAF1 | GTP-RAS-bound RAF1 |
| Ras_iRAF1 | GTP-RAS-bound inhibited RAF1 |
| Ras_BRAF | GTP-RAS-bound BRAF |
| Ras_iBRAF | GTP-RAS-bound inhibited BRAF |
| Ras_pRaf1 | GTP-RAS-bound phosphorylated RAF1 |
| Ras_pRaf1_Raf1_tetramer | Homodimerized active RAF1 and phosphorylated RAF1 |
| Ras_pRaf1_iRaf1_tetramer | Homodimerized inhibited RAF1 and phosphorylated RAF1 |
| Ras_Braf_Raf1_tetramer | Heterodimerized active BRAF and active RAF1 |
| Ras_iRaf1_tetramer | GTP-Ras-bound inhibited RAF1 |
| Ras_Braf_iRaf1_tetramer | Heterodimerized active BRAF and inhibited RAF1 |
| Ras_pRaf1_tetramer | Phosphorylated pRAF1-RAF1 homodimer |
| Ras_Braf_pRaf1_tetramer | Phosphorylated BRAF-RAF1 heterodimer |
| Ras_Raf1_iBraf_tetramer | GTP-RAS-bound RAF1-inhibited BRAF dimer |
| Ras_pRaf1_iBraf_tetramer | Phosphorylated iBRAF-RAF1 heterodimer |
| Ras_iRaf1_iBraf1_tetramer | Inhibited RAF1-inhibited BRAF dimer |
| MEK | MEK1/2 |
| pMEK | Phosphorylated MEK1/2 |
| ERK | ERK1/2 |
| pERK | EGF-bound, GRB2-pGAB1-SHP2-bound pEGFR dimer |
| nfpSOS | negative feedback phosphorylated SOS |
| nfpRaf1 | negative feedback phosphorylated RAF1 |
| nfpIRaf1 | negative feedback phosphorylated inhibited RAF 1 |
| nfpBRaf | negative feedback phosphorylated BRAF |
| nfpIBRaf | negative feedback phosphorylated inhibited BRAF |
| Ras_nfpRaf1 | GTP-RAS-bound negative feedback phosphorylated RAF1 |
| Ras_nfpBRaf | GTP-RAS-bound negative feedback phosphorylated BRAF |
| Ras_nfpiBRaf | GTP-RAS-bound negative feedback phosphorylated inhibited BRAF |
| Ras_nfpiRaf1 | GTP-RAS-bound negative feedback phosphorylated inhibited RAF1 |
| sorafenib | RAF1/BRAF inhibitor |

**Table S4.** Best-fit values of rate constants from SloppyCell and initial conditions used in the EGFR-ERK model analyses.

|  | Parameter (units) | Description | Value | Source |
| --- | --- | --- | --- | --- |
| 1 | $k_{Ef}$ ($\mu\text{M}^{-1} \text{min}^{-1}$) | EGF binding to EGFR, forward | $1.251 \times 10^4$ | Fit |
| 2 | $k_{Er}$ ($\text{min}^{-1}$) | EGF binding to EGFR, reverse | $2.157 \times 10$ | Fit |
| 3 | $k_{dEf}$ ($\text{cell molec}^{-1} \text{min}^{-1}$) | EGFR dimerization, forward | $9.793 \times 10^{-1}$ | Fit |
| 4 | $k_{dEr}$ ($\text{min}^{-1}$) | EGFR dimerization, reverse | $1.518 \times 10$ | Fit |
| 5 | $k_{catE}$ ($\text{min}^{-1}$) | EGFR phosphorylation, EGF-occupied dimer | 6.663 | Fit |
| 6 | $k_{dp}$ ($\text{min}^{-1}$) | EGFR dephosphorylation | $5.178 \times 10$ | Fit |
| 7 | $k_{G2f}$ ($\text{cell molec}^{-1} \text{min}^{-1}$) | GRB2 binding to EGFR, forward | $6.520 \times 10^{-1}$ | Fit |
| 8 | $k_{G2r}$ ($\text{min}^{-1}$) | GRB2 binding to EGFR, reverse | $3.334 \times 10^3$ | Fit |
| 9 | $k_{SOSf}$ ($\text{cell molec}^{-1} \text{min}^{-1}$) | SOS binding to GRB2, forward<br>EGFR-GRB2 binding to SOS, forward | $5.520 \times 10^{-5}$ | Fit |
| 10 | $k_{SOSr}$ ($\text{min}^{-1}$) | SOS binding to GRB2, reverse<br>EGFR-GRB2 binding to SOS, reverse | 1.905 | Fit |
| 11 | $k_{nfpSOS}$ ($\text{cell molec}^{-1} \text{min}^{-1}$) | Feedback phosphorylation of SOS by ERK | $7.836 \times 10^{-6}$ | Fit |
| 12 | $k_{dpSOS}$ ($\text{min}^{-1}$) | Dephosphorylation of feedback-phosphorylated SOS | $5.861 \times 10^{-3}$ | Fit |
| 13 | $k_{Rgneslow}$ ($\text{min}^{-1}$) | SOS catalyzed GDP-to-GTP exchange on Ras | 5.755 | Fit |
| 14 | $K_{mgneslow}$ ($\text{molec cell}^{-1}$) | Michaelis constant for SOS-catalyzed GDP-to-GTP exchange on RAS | $7.205 \times 10^4$ | Fit |
| 15 | $k_{Rhydro}$ ($\text{min}^{-1}$) | Hydrolyzation of Ras-bound GTP | $2.712 \times 10^{-1}$ | Fit |
| 16 | $k_{R1f}$ ($\text{cell molec}^{-1} \text{min}^{-1}$) | RAF1 binding to RAS-GTP, forward | $3.314 \times 10^{-5}$ | Fit |
| 17 | $k_{R1r}$ ($\text{min}^{-1}$) | RAF1 binding to RAS-GTP, reverse | $9.084 \times 10$ | Fit |
| 18 | $k_{Bf}$ ($\text{cell molec}^{-1} \text{min}^{-1}$) | BRAF binding to RAS-GTP, forward | $5.634 \times 10^2$ | Fit |
| 19 | $k_{Br}$ ($\text{min}^{-1}$) | BRAF binding to RAS-GTP, reverse | $1.726 \times 10^{-2}$ | Fit |
| 20 | $k_{fpR1r}$ ($\text{min}^{-1}$) | Feedback-phosphorylated RAF1 unbinding from RAS-GTP, forward | $4.116 \times 10^{-2}$ | Fit |
| 21 | $k_{fpBr}$ ($\text{min}^{-1}$) | Feedback-phosphorylated BRAF unbinding from RAS-GTP, forward | 5.323 | Fit |
| 22 | $k_{dR1f}$ ($\text{cell molec}^{-1} \text{min}^{-1}$) | RAF1 dimerization, forward | $9.754 \times 10^{-1}$ | Fit |
| 23 | $k_{dR1r}$ ($\text{min}^{-1}$) | RAF1 dimerization, reverse | $1.314 \times 10$ | Fit |
| 24 | $k_{pR1}$ ($\text{min}^{-1}$) | RAF1 phosphorylation | $2.289 \times 10^3$ | Fit |
| 25 | $k_{dpR1}$ ($\text{min}^{-1}$) | RAF1 dephosphorylation | $2.813 \times 10$ | Fit |
| 26 | $k_{nfpBR1}$ ($\text{cell molec}^{-1} \text{min}^{-1}$) | Feedback phosphorylation of RAF1, BRAF, iRAF1, and iBRAF by ERK | $3.806 \times 10^{-13}$ | Fit |
| 27 | $k_{\text{Son}} (\mu\text{M}^{-1} \text{min}^{-1})$ | Sorafenib binding to RAF1/BRAF, forward | $2.828 \times 10$ | Fit |
| 28 | $k_{\text{Soff}} (\text{min}^{-1})$ | Sorafenib binding to RAF1/BRAF, reverse | 8.723 | Fit |
| 29 | $k_{\text{iR1f}} (\text{cell molec}^{-1} \text{min}^{-1})$ | Inhibited RAF1 binding to RAS-GTP, forward | $1.480 \times 10^{-6}$ | Fit |
| 30 | $k_{\text{iR1r}} (\text{min}^{-1})$ | Inhibited RAF1 binding to RAS-GTP, reverse | $2.202 \times 10^{-1}$ | Fit |
| 31 | $k_{\text{pMEK}} (\text{cell molec}^{-1} \text{min}^{-1})$ | MEK phosphorylation | $1.588 \times 10^{-4}$ | Fit |
| 32 | $k_{\text{dpMEK}} (\text{min}^{-1})$ | MEK dephosphorylation | $5.110 \times 10^{-1}$ | Fit |
| 33 | $k_{\text{pERK}} (\text{cell molec}^{-1} \text{min}^{-1})$ | ERK phosphorylation | $1.161 \times 10^{-6}$ | Fit |
| 34 | $k_{\text{dpERK}} (\text{min}^{-1})$ | ERK dephosphorylation | $3.766 \times 10^{-1}$ | Fit |
| 35 | $k_{\text{G2SOSf}} (\text{cell molec}^{-1} \text{min}^{-1})$ | EGF-occupied EGFR binding to GRB2-SOS | $2.594 \times 10^{-1}$ | Fit |
| 36 | $k_{\text{G2SOSr}} (\text{min}^{-1})$ | EGF-occupied EGFR unbinding from GRB2-SOS | $5.888 \times 10$ | Fit |
| 37 | $k_{\text{nfpiR1r}} (\text{min}^{-1})$ | Feedback-phosphorylated iRAF1 unbinding from RAS-GTP, reverse | $1.742 \times 10^{-2}$ | Fit |
| 38 | $k_{\text{nfpiBr}} (\text{min}^{-1})$ | Feedback-phosphorylated iBRAF unbinding from RAS-GTP, reverse | $1.022 \times 10$ | Fit |
| 39 | $k_{\text{iBf}} (\text{cell molec}^{-1} \text{min}^{-1})$ | iBRAF binding to RAS-GTP, forward | $1.580 \times 10^{-13}$ | Fit |
| 40 | $k_{\text{iBr}} (\text{min}^{-1})$ | iBRAF binding to RAS-GTP, reverse | 1.564 | Fit |
| 41 | Sorafenib ($\mu\text{M}$) | Sorafenib concentration | 10 | Surve *et al.*, 2019 |
| 42 | EGF ($\mu\text{M}$) | Extracellular EGF concentration | $1.7 \times 10^{-3}$ | Surve *et al.*, 2019 |
| 43 | EGFR ($\text{cell}^{-1}$) | EGFR molecules per HeLa cell | 93,000 | Kulak *et al.*, 2014 |
| 44 | GRB2 ($\text{cell}^{-1}$) | GRB2 molecules per HeLa cell | 628,000 | Kulak *et al.*, 2014 |
| 45 | SOS ($\text{cell}^{-1}$) | SOS1/2 molecules per HeLa cell | 7700 | Kulak *et al.*, 2014 |
| 46 | RAF1 ($\text{cell}^{-1}$) | RAF1 molecules per HeLa cell | 12,000 | Shi *et al.*, 2016 |
| 47 | BRAF ($\text{cell}^{-1}$) | BRAF molecules per HeLa cell | 1000 | Kulak *et al.*, 2014 |
| 48 | RAS ($\text{cell}^{-1}$) | RAS molecules per HeLa cell (HRAS, NRAS, KRAS) | 135,000 | Kulak *et al.*, 2014 |
| 49 | MEK ($\text{cell}^{-1}$) | MEK1/2 molecules per HeLa cell | 500,000 | Kulak *et al.*, 2014 |
| 50 | ERK ($\text{cell}^{-1}$) | ERK1/2 molecules per HeLa cell | 600,000 | Kulak *et al.*, 2014 |

**Table S5.** Cost for model fitting using the VIP > 1 parameters identified from PLSR SA versus all parameters in SloppyCell.

| Model fitting method | Cost ($C(\theta)$) |
| --- | --- |
| PLSR SA | 466.08 |
| Sloppiness Analysis | 123.86 |

**Table S6.** Comparison of cost, computational time, and residual error of single best model fits by varying different numbers of parameters of importance from PLSR SA.

| Number of parameters allowed to vary in fitting | SA sampling method | VIP score threshold | Wall-clock time (s) | Time (min) | Residual sum of squares | Max. iterations allowed |
| --- | --- | --- | --- | --- | --- | --- |
| 40 | - | - | 2405.88 | 40.10 | 468.51 | 10000 |
| 22 | Log-uniform random | 0.5 | 1237.44 | 20.62 | 469.02 | 10000 |
| 20 | Log-uniform LHS | 0.5 | 1570.27 | 26.17 | 462.77 | 10000 |
| 15 | Log-uniform random | 0.8 | 706.33 | 11.77 | 474.84 | 10000 |
| 15 | Log-uniform LHS | 0.8 | 658.20 | 10.97 | 474.84 | 10000 |
| 9 | Log-uniform random | 1 | 264.35 | 4.41 | 619.70 | 10000 |
| 10 | Log-uniform LHS | 1 | 450.07 | 7.50 | 607.69 | 10000 |

**Table S7.** Comparison of the best fit parameter values when a subset of important parameters were allowed to vary vs. all parameters were allowed to vary.

|  | Parameter | VIP > 1<br>parameters varied | All (40)<br>parameters varied | Reference<br>values | Source |
| --- | --- | --- | --- | --- | --- |
| 1 | K<sub>Ef</sub> | | $1.4 \times 10^4$ | $3.1 \times 10^2$ | Waters *et al.*, 1990 |
| 2 | K<sub>catE</sub> | | $1.6 \times 10$ | $1.3 \times 10$ | Fan *et al.*, 2004 |
| 3 | K<sub>pMEK</sub> | $3.6 \times 10^{-4}$ | $1.6 \times 10^{-4}$ | $6.0 \times 10^{-6}$ | Varga *et al.*, 2017 |
| 4 | K<sub>nfpSOS</sub> | $2.1 \times 10^{-4}$ | $1.0 \times 10^{-5}$ | $2.5 \times 10^{-3}$ | Surve *et al.*, 2019 |
| 5 | K<sub>Br</sub> | | $2.1 \times 10^{-2}$ | $3.7 \times 10^{-2}$ | Fischer *et al.*, 2007 |
| 6 | K<sub>Bf</sub> | | $5.0 \times 10^2$ | $2.5 \times 10^{-7}$ | Fischer *et al.*, 2007 |
| 7 | K<sub>dpERK</sub> | 1.9 | $4.5 \times 10^{-1}$ | $6.0 \times 10^{-1}$ | Chen *et al.*, 2009 |
| 8 | K<sub>R1r</sub> | $1.4 \times 10$ | $8.1 \times 10$ | 8.4 | Surve *et al.*, 2019 |
| 9 | K<sub>R1f</sub> | | $3.6 \times 10^{-5}$ | $2.5 \times 10^{-5}$ | Surve *et al.*, 2019 |
| 10 | K<sub>nfpBR1</sub> | | $1.0 \times 10^{-6}$ | $3.7 \times 10^{-1}$ | Varga *et al.*, 2017 |
| 11 | K<sub>pR1</sub> | | $3.7 \times 10^3$ | $1.9 \times 10^2$ | Chen *et al.*, 2009 |
| 12 | K<sub>SOSr</sub> | | 2.2 | $8.3 \times 10^{-3}$ | Surve *et al.*, 2019 |
| 13 | K<sub>SOSf</sub> | | $6.1 \times 10^{-5}$ | $2.2 \times 10^{-6}$ | Surve *et al.*, 2019 |
| 14 | K<sub>Rgneslow</sub> | $1.7 \times 10^1$ | 5.9 | $1.8 \times 10^{-1}$ | Das *et al.*, 2009 |
| 15 | K<sub>Son</sub> | | $3.1 \times 10$ | $1.0 \times 10^5$ | Surve *et al.*, 2019 |
| 16 | K<sub>G2SOSr</sub> | | $4.9 \times 10$ | $8.3 \times 10^{-3}$ | Kholodenko *et al.*, 1999 |
| 17 | K<sub>G2SOSf</sub> | $2.4 \times 10^{-2}$ | $2.0 \times 10^{-1}$ | $2.2 \times 10^{-6}$ | Kholodenko *et al.*, 1999 |
| 18 | K<sub>iR1r</sub> | | $2.3 \times 10^{-1}$ | $1.4 \times 10^{-2}$ | Surve *et al.*, 2019 |
| 19 | K<sub>iR1f</sub> | $2.1 \times 10^{-5}$ | $1.0 \times 10^{-6}$ | $1.1 \times 10^{-6}$ | Surve *et al.*, 2019 |
| 20 | K<sub>Soff</sub> | | $7.2 \times 10^{-1}$ | $4.2 \times 10^2$ | Surve *et al.*, 2019 |
| 21 | K<sub>pERK</sub> | $7.0 \times 10^{-6}$ | $1.0 \times 10^{-6}$ | $6.0 \times 10^{-5}$ | Varga *et al.*, 2017 |
| 22 | K<sub>Rhydro</sub> | $2.4 \times 10^{-1}$ | $3.2 \times 10^{-1}$ | $1.6 \times 10^{-1}$ | Das *et al.*, 2009 |
| 23 | K<sub>mgneslow</sub> | $6.8 \times 10^5$ | $7.3 \times 10^4$ | $1.4 \times 10^5$ | Das *et al.*, 2009 |
| 24 | K<sub>dpMEK</sub> | $2.4 \times 10^{-2}$ | $5.4 \times 10^{-1}$ | $6.0 \times 10^{-1}$ | Chen *et al.*, 2009 |
| 25 | K<sub>nfpIR1r</sub> | | $1.7 \times 10^{-2}$ | $4.6 \times 10^{-2}$ | Surve *et al.*, 2019 |
| 26 | K<sub>dR1r</sub> | | $1.9 \times 10$ | $1.1 \times 10^2$ | Surve *et al.*, 2019 |
| 27 | K<sub>iBr</sub> | | 1.2 | $3.7 \times 10^{-2}$ | Surve *et al.*, 2019 |
| 28 | K<sub>dR1f</sub> | | 1.2 | $1.9 \times 10^{-2}$ | Surve *et al.*, 2019 |
| 29 | K<sub>iBf</sub> | | $1.0 \times 10^{-6}$ | $2.5 \times 10^{-7}$ | Surve *et al.*, 2019 |
| 30 | K<sub>fpBr</sub> | | 4.3 | $3.7 \times 10^{-1}$ | Surve *et al.*, 2019 |
| 31 | K<sub>dpSOS</sub> | | $2.3 \times 10^{-4}$ | $6.0 \times 10^{-1}$ | Surve *et al.*, 2019 |
| 32 | K<sub>dp</sub> | | $2.7 \times 10$ | 8 | Furcht *et al.*, 2015 |
| 33 | K<sub>dEr</sub> | | $2.5 \times 10$ | $1.0 \times 10^{-1}$ | Hendriks *et al.*, 2003 |
| 34 | K<sub>dEf</sub> | | $6.1 \times 10^{-1}$ | $9.2 \times 10^{-4}$ | Furcht *et al.*, 2015 |
| 35 | K<sub>dpR1</sub> | | $4.2 \times 10$ | $6.0 \times 10^{-1}$ | Chen *et al.*, 2009 |
| 36 | K<sub>G2r</sub> | | $3.3 \times 10^3$ | $3.8 \times 10^{-4}$ | Morimatsu *et al.*, 2007 |
| 37 | K<sub>G2f</sub> | | $5.5 \times 10^{-1}$ | $4.6 \times 10^2$ | Morimatsu *et al.*, 2007 |
| 38 | $k_{\text{nfpiBr}}$ | | 8.3 | $3.7 \times 10^{-1}$ | Surve *et al.*, 2019 |
| 39 | $k_{\text{fpR1r}}$ | | $2.3 \times 10^{-2}$ | 2.1 | Surve *et al.*, 2019 |
| 40 | $k_{\text{Er}}$ | | $2.9 \times 10$ | $8.0 \times 10^{-1}$ | French *et al.*, 1995 |

